# Molecular tension authenticates apoptotic cells being phagocytosed

**DOI:** 10.1101/2024.06.10.598189

**Authors:** Chanhyuk Min, Hyeokjin Cho, Jeongmin Yu, Byeongjin Moon, Jaeseon Jeon, Nafeesa Shahdab, Hyunji Moon, Susumin Yang, Juyeon Lee, Jaeseong Jin, Mingi Hong, Won-jin Chung, Jihwan Park, Gwangrog Lee, Daeho Park

**Author notes:** Correspondence and requests for materials should be addressed to: G.L. or D.P. These authors contributed equally: Chanhyuk Min, Hyeokjin Cho, Jeongmin Yu. School of Health and Life Sciences, Teesside University; Middlesbrough, United Kingdom (Nafeesa Shahdab).

## Abstract

Profound cytoskeletal reorganization and plasma membrane deformation in phagocytes is indispensable for phagocytosis of massive apoptotic cells, but whether these dynamics accompany a mechanical signal modulating signaling during efferocytosis remains largely unexplored. Here, we report that tension between phosphatidylserine (PS) and PS receptors (PSR) generated at the phagocytic synapse serves as a signal to determine whether PS-exposing cells should be phagocytosed. Mechanistically, increased membrane tension of phagocytes via Rac1-dependent actin polymerization and membrane stiffness of apoptotic cells caused tension between PS and PSRs, leading to phosphoinositide 3-kinase recruitment to PSRs, which resulted in Rac1 inactivation and myosin II phosphorylation required for phagocytic cup closure. Our observations imply that tension between PS and PSRs acts as a mechanical fail-safe to prevent removal of all PS-exposing cells for the integrity of efferocytosis.

**One-Sentence Summary:** Tension between phosphatidylserine and its receptors acts as a final decision-maker determining phagocytosis of apoptotic cells.

## Main Text

Phagocytosis of apoptotic cells, called efferocytosis, differs from other types of phagocytosis in terms of the endogeny and size of the targets. Specifically, this process removes endogenously generated apoptotic cells that are as large as phagocytes (*1, 2*). Elaborately coordinated signaling between two types of biochemical signals, called ‘eat-me’ and ‘don’t eat-me’ signals, enables phagocytes to remove such endogenous and massive targets. Phagocytes sense cells to be phagocytosed through eat-me signals, of which phosphatidylserine (PS) is the most well-known, and cells not to be phagocytosed through don’t eat-me signals (*3–8*). Accordingly, when eat-me signals are present and don’t eat-me signals are nullified on the cell surface, phagocytes are convinced that the cells are apoptotic cells that should be phagocytosed, and ultimately efferocytosis may occur (*9, 10*). Furthermore, concerted signaling between these two signals induces cytoskeletal rearrangement, which results in plasma membrane deformation, allowing phagocytes to engulf the massive targets (*11–13*). Nevertheless, it has not been sufficiently assessed whether the two signals are decisive for efferocytosis to occur. In particular, the drastic plasma membrane deformation and cytoskeletal rearrangement in phagocytes may induce a tensile force (tension) between molecules interacting intra- and intercellularly on the plasma membrane (e.g., PS on apoptotic cells and its receptors (PSRs) on phagocytes). This force may interconnect with the conventional signaling of efferocytosis. However, it remains unclear whether such a force exists and, if so, what role it plays in efferocytosis. In this study, we investigated the existence, generation, and role of tension between PS and PSRs (referred to as tension of PS-PSRs) in efferocytosis using various approaches including surrogates of apoptotic cells that can be used to adjust tension of PS-PSRs and time-lapse confocal microscopy.

To investigate whether tension of PS-PSRs exists during and is required for efferocytosis, we devised beads with PS connected via a tension gauge tether (*14*) (PS-TGT) as surrogates of apoptotic cells. In PS-TGT beads, PS is covalently conjugated to the Cy3- or Cy5-labeled upper strand of a double-stranded DNA tether that is immobilized on neutravidin beads and the TGT ruptures at a critical force, termed tension tolerance (Fig. 1A). When J774A.1 cells, a murine macrophage cell line, were incubated with PS-TGT beads, the cells engulfed beads with a 56 pN tension tolerance TGT (PS-TGT^56pN^) more efficiently than beads with a 12 pN tension tolerance TGT (PS-TGT^12pN^), as measured by the number of engulfed beads per phagocyte and the percentage of phagocytes engulfing the beads (Fig. 1, B and C, and movie S1). Other macrophages, including peritoneal macrophages, bone marrow-derived macrophages, and RAW264.7 cells, also engulfed PS-TGT^56pN^ beads more efficiently than PS-TGT^12pN^ beads (fig. S1, A to C). However, the superior engulfment of TGT^56pN^ beads relative to TGT^12pN^ beads by macrophages was not observed when phosphatidylcholine (PC) was conjugated to the TGT instead of PS (Fig. 1D). Notably, these results were not dependent on the amount of TGT immobilized on the beads because PS-TGT^56pN^ beads were consistently engulfed more efficiently than PS-TGT^12pN^ beads and engulfment of PC-TGT^56pN^ beads was still comparable with that of PC-TGT^12pN^ beads when various amounts of the TGT were immobilized on the beads (fig. S1D).

**Fig. 1.**
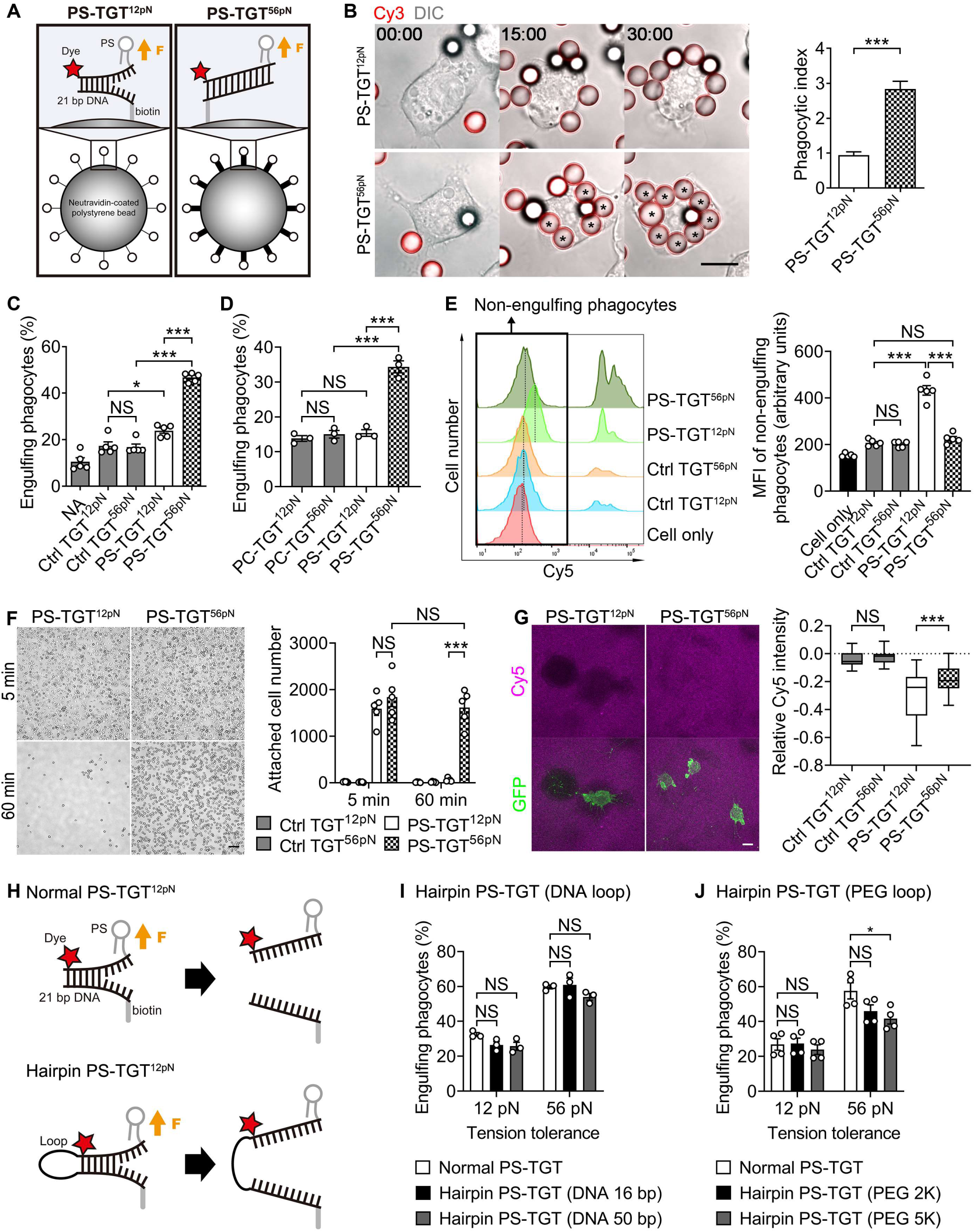
Tension-dependent engulfment of PS-TGT beads. (**A**) Schematic diagram of PS-TGT beads. PS and biotin were conjugated to the Cy3- or Cy5-labeled upper strand and the lower strand of 21 bp DNA, respectively, which was immobilized on neutravidin-coated polystyrene beads via a neutravidin-biotin linker. Yellow arrows indicate the direction of force (F). (**B**) J774A.1 cells incubated with Cy3-labeled PS-TGT^12pN^ or PS-TGT^56pN^ beads for 30 min were observed by time-lapse confocal microscopy (left) and engulfed beads were counted (right). Asterisks indicate ingested beads. Scale bar, 10 μm. Data are mean ± s.e.m. from n=165 cells for PS-TGT^12pN^ and n=124 cells for PS-TGT^56pN^. ***P < 0.001; two-tailed unpaired Student’s *t* test. (**C** and **D**) J774A.1 cells were incubated with the indicated beads for 10 min and analyzed by flow cytometry. Ctrl, control; NA, neutravidin bead. Data are mean ± s.e.m. from n=5 (C) and n=3 (D) independent experiments. *P < 0.05; ***P < 0.001; NS, not significant; one-way ANOVA. (**E**) J774A.1 cells incubated with the indicated beads for 10 min were analyzed by flow cytometry. Representative histograms of phagocytes (left) and MFIs of non-engulfing phagocytes (right) are shown. The box and dotted lines in the histograms indicate non-engulfing phagocytes and the peak of the fluorescence intensity of non-engulfing phagocytes, respectively. Data are mean ± s.e.m. from n=5 independent experiments. ***P < 0.001; NS, not significant; one-way ANOVA. (**F**) J774A.1 cells were incubated on PS-TGT plates, on which the indicated PS-TGTs were immobilized, for 5 min, washed, further incubated for 60 min, and washed to remove detached cells. Scale bar, 200 μm. Data are mean ± s.e.m. from n=5 independent experiments. ***P < 0.001; NS, not significant; two-way ANOVA. (**G**) J774A.1 cells stably expressing membrane-targeted GFP (GFP-CAAX) were incubated on PS-TGT plates, on which a mixture of the indicated Cy5-labeled PS-TGTs and Cy3-labeled PS-TGT^56pN^ was immobilized, for 60 min and observed by confocal microscopy. Relative Cy5 intensities were calculated. The intensity of Cy5 in the area where cells were absent was set to 0. Scale bar, 10 μm. Data are represented in the box-and-whisker plot. n=26 cells for Ctrl TGT^12pN^, n=30 cells for Ctrl TGT^56pN^, n=29 cells for PS-TGT^12pN^, and n=49 cells for PS-TGT^56pN^. ***P < 0.001; NS, not significant; one-way ANOVA. (**H**) Schematic diagram of normal PS-TGT^12pN^ (top) and hairpin PS-TGT^12pN^ (bottom). The upper and lower strands of hairpin PS-TGT^12pN^ were connected by PEG or DNA loops. Yellow arrows indicate the direction of force (F). (**I** and **J**) J774A.1 cells were incubated with the indicated beads for 15 min. Percentage of engulfing phagocytes were measured by flow cytometry. Data are mean ± s.e.m. from n=3 independent experiments for (I), n=4 independent experiments for (J). *P < 0.05; NS, not significant; two-way ANOVA.

We then investigated whether the TGT is indeed ruptured during engulfment of PS-TGT beads. Several observations indicated that PS-TGT^12pN^, but not PS-TGT^56pN^, was ruptured during engulfment of the beads. First, the Cy5 mean fluorescence intensity (MFI) of non-engulfing phagocytes was higher upon incubation with PS-TGT^12pN^ beads than upon incubation with PS-TGT^56pN^ or control TGT beads (Fig. 1E), implying that the ruptured Cy5-labeled upper strand of PS-TGT^12pN^ is present in non-engulfing phagocytes. Second, after macrophages were incubated on a polyethylene glycol (PEG)-passivated glass plate on which PS-TGT was immobilized via a biotin-neutravidin interaction (*15*), cell detachment from the plate was evaluated. The cell number was comparable on the PS-TGT^12pN^ and PS-TGT^56pN^ plates after incubation for 5 min. However, after incubation for 60 min, few cells were present on the PS-TGT^12pN^ plate, whereas most cells remained attached to the PS-TGT^56pN^ plate (Fig. 1F). This also implies that PS-TGT^12pN^, but not PS-TGT^56pN^, is ruptured because the cells were attached to the plate only through PS-TGT (fig. S1E). Third, the loss of Cy5 fluorescence, indicative of rupture of the Cy5-labeled upper strand of the TGT, was greater with PS-TGT^12pN^ than with PS-TGT^56pN^ when macrophages were incubated on glass plates (Fig. 1G and fig. S1F). Collectively, these data imply that the degree of rupture of the PS-TGT differs depending on tension tolerance during engulfment of PS-TGT beads, indicating that tension is loaded between PS and PSRs during and is essential for efferocytosis.

The rupture of PS-TGT^12pN^ may cause the ligand to separate from the target, preventing further signaling and potentially leading to inefficient engulfment of the PS-TGT^12pN^ beads. To test this, we designed hairpin PS-TGTs with variable loop sizes, created using DNA or PEG, that prevent the ligand from separating from the phagocytic target even when TGT rupture occurs. In this measurement, the upper and lower strands of 12 pN TGT were connected by loops made of PEG or ssDNA, creating TGTs with various tensions depending on the length of the loop (Fig. 1H). For the hairpin TGTs with ssDNA after rupture (Fig. 1I), they are subjected to minimal entropic tension of ∼2.3 and ∼3.6 pN for the 16-bp- and 50-bp-ssDNA loops, respectively, based on the length-dependent elasticity of ssDNA. The results showed that macrophages still engulfed the hairpin PS-TGT^12pN^ beads with significantly lower efficiency than hairpin or normal PS-TGT^56pN^ beads, and there were no differences between engulfment of normal PS-TGT^12pN^ and hairpin PS-TGT^12pN^ beads. Nevertheless, in the case of the PEG loop (Fig. 1J), as the loop size increased, engulfment decreased at both 12 pN and 56 pN, which is believed to be due to increased steric hindrance as loop size increases. Regardless of the loop size and type, the engulfment of hairpin PS-TGT^12pN^ beads remained less efficient compared to the engulfment of hairpin PS-TGT^56pN^ beads (Fig. 1, I and J), implying that inefficient engulfment of PS-TGT^12pN^ beads is not due to the termination of signaling caused by the disassembly of the ligand from the phagocytic target.

We next investigated how tension of PS-PSRs affects engulfment of PS-TGT beads. To this end, we scrutinized the binding, internalization, and degradation steps of efferocytosis. The numbers of PS-TGT^12pN^ and PS-TGT^56pN^ beads bound to macrophages were comparable when cells were incubated with the beads at 4°C (fig. S2A). F-actin formation, which is required for internalization of apoptotic cells during efferocytosis and was monitored by phalloidin staining, was also comparable around PS-TGT^12pN^ and PS-TGT^56pN^ beads (fig. S2B). In addition, neither the acidification of phagolysosomes nor the degradation of phagolysosomal cargos, as measured by the change rates of the fluorescence intensities of pHrodo and Cy3, respectively, was altered according to the tension tolerance of PS-TGT beads (fig. S2, C and D, and movies S2 and S3). However, unexpectedly, time-lapse observation of F-actin using SiR-actin, a live F-actin labeling probe, revealed that F-actin persisted longer around PS-TGT^12pN^ beads than around PS-TGT^56pN^ beads (Fig. 2, A and D, and movie S4). Disassembly of F-actin is synchronized with phagocytic cup closure during efferocytosis (*16, 17*); therefore, we next monitored phagocytic cup closure depending on tension tolerance during engulfment of PS-TGT beads. Macrophages often failed to close the phagocytic cup to completely engulf PS-TGT^12pN^ beads and consequently it took longer to close the phagocytic cup for engulfment of PS-TGT^12pN^ beads than for engulfment of PS-TGT^56pN^ beads (Fig. 2, C and D, and movie S5). These data imply that tension of PS-PSRs affects efferocytosis by modulating phagocytic cup closure but not by modulating the binding capacity, F-actin formation, or phagolysosomal function of phagocytes during efferocytosis.

**Fig. 2.**
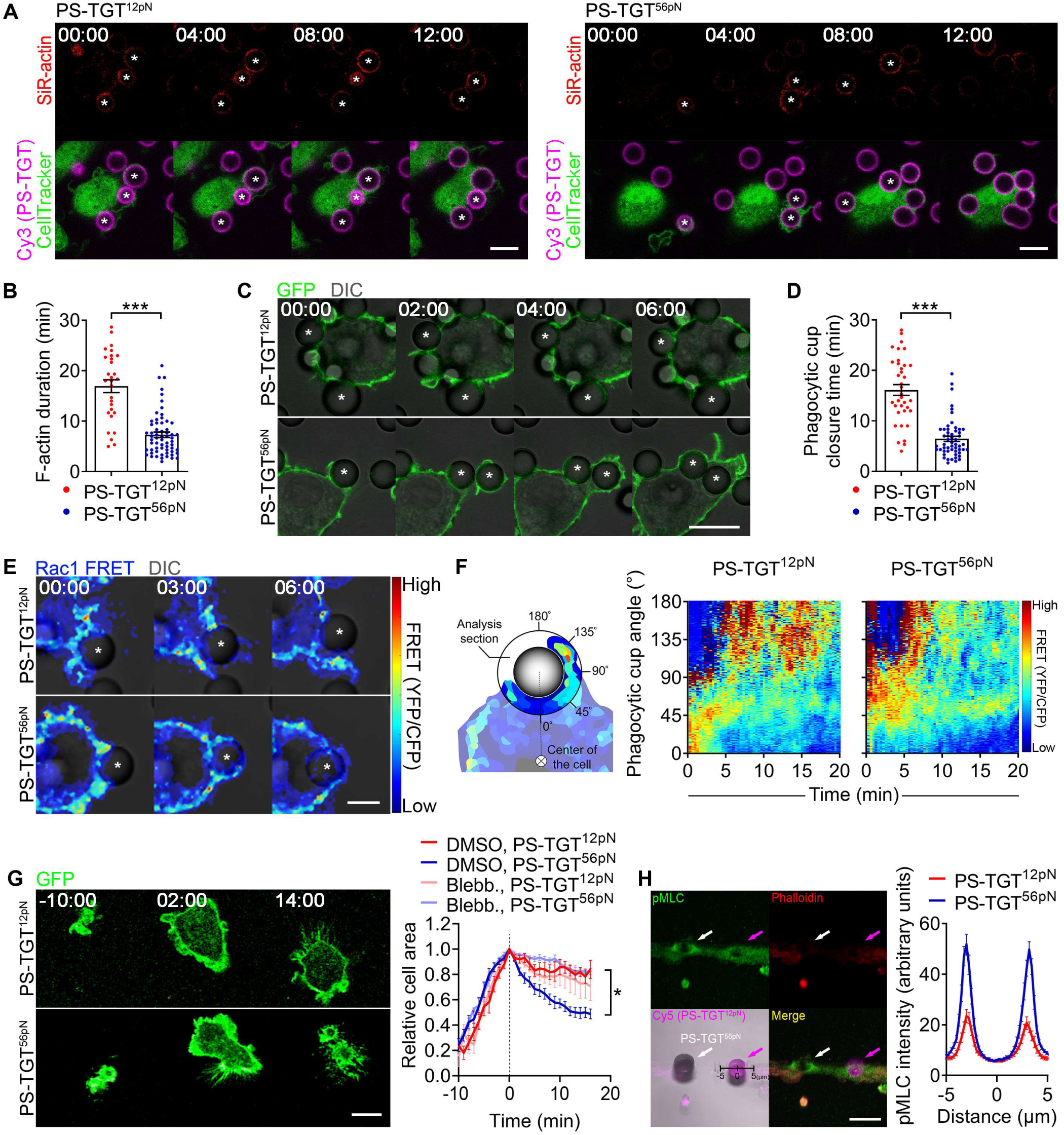
The tension affects engulfment of PS-TGT beads through Rac1. (**A** and **B**) J774A.1 cells stained with SiR-actin were incubated with Cy3-labeled PS-TGT beads for 30 min and observed by time-lapse confocal microscopy (A). The duration of F-actin labeling was measured (B). Asterisks indicate PS-TGT beads with SiR-actin signals. Scale bar, 10 μm. Data are mean ± s.e.m. from n=29 beads for PS-TGT^12pN^ and n=62 beads for PS-TGT^56pN^. ***P < 0.001; two-tailed unpaired Student’s *t* test. (**C** and **D**) J774A.1 cells stably expressing membrane-targeted GFP were incubated with Cy3-labeled PS-TGT beads for 30 min and observed by time-lapse confocal microscopy (C). Asterisks indicate beads with extended phagocytic cups. The amount of time required for phagocytic cup closure was measured (D). Scale bar, 10 μm. Data are mean ± s.e.m. from n=36 beads for PS-TGT^12pN^ and n=54 beads for PS-TGT^56pN^. ***P < 0.001; two-tailed unpaired Student’s *t* test. (**E** and **F**) J774A.1 cells stably expressing Raichu-Rac1 were incubated with PS-TGT beads for 30 min and observed by time-lapse confocal microscopy (E). Asterisks indicate beads with extended phagocytic cups. Rac1 activity in phagocytic cups around the beads was calculated by YFP/CFP ratio FRET. Scale bar, 5 μm. Schematic diagram depicting the analysis of Rac1 FRET according to the phagocytic cup position (F, left). Rac1 activity was measured at the phagocytic cup around the beads (F, right). n=33 beads for PS-TGT^12pN^ and n=23 beads for PS-TGT^56pN^. (**G**) J774A.1 cells stably expressing membrane-targeted GFP were incubated on the indicated PS-TGT plates with or without blebbistatin (10 μM) and observed by time-lapse confocal microscopy. When the cells were maximally expanded, the time and relative area were set to 0 min and 1, respectively. The areas of cells on the plates were normalized to the maximum area. Scale bar, 10 μm. Blebb, blebbistatin. Data are mean ± s.e.m. from n=8 cells for PS-TGT^12pN^ (DMSO), n=11 cells for PS-TGT^56pN^ (DMSO), n=16 cells for PS-TGT^12pN^ (Blebb.), and n=8 cells for PS-TGT^56pN^ (Blebb.). Statistical analysis was performed between the end points of PS-TGT^12pN^ (DMSO) and PS-TGT^56pN^ (DMSO). *P < 0.05; one-way ANOVA. (**H**) J774A.1 cells were incubated with both Cy5-labeled PS-TGT^12pN^ and unlabeled PS-TGT^56pN^ beads for 10 min, stained with an anti-phospho-MLC antibody and Alexa Fluor 594-conjugated phalloidin, and observed by confocal microscopy. Magenta and white arrows indicate PS-TGT^12pN^ and PS-TGT^56pN^ beads, respectively. The intensities of phosphorylated MLC (pMLC) staining were quantified across the phagocytic cups. Scale bar, 10 μm. Data are mean ± s.e.m. from n=26 beads for PS-TGT^12pN^ and PS-TGT^56pN^.

We then investigated how tension of PS-PSRs modulates phagocytic cup closure during engulfment of PS-TGT beads. Rac1 is involved in an early signaling pathway leading to efferocytosis and its activity regulates F-actin dynamics (*11, 18*). In particular, spatiotemporal regulation of Rac1 activity at the phagocytic cup is crucial for phagocytic cup formation and closure (*16*). Therefore, we investigated whether the tension affects Rac1 activity. To this end, we used Raichu-Rac1, a biosensor that monitors Rac1 activity via fluorescence resonance energy transfer (FRET) (*19*). Macrophages expressing Raichu-Rac1 were incubated with PS-TGT beads, and FRET was measured at the interface between macrophages and the beads from 0° to 180° in a circular fashion as a function of time. Rac1 activation was initiated at the junction of macrophages and the beads and proceeded along the tip of the phagocytic cup. At 0–10 min, Rac1 was activated with similar FRET levels in phagocytes incubated with PS-TGT^12pN^ and PS-TGT^56pN^ beads. By contrast, at 10–20 min, Rac1 activation was instantly followed by its inactivation at the phagocytic cup of phagocytes incubated with PS-TGT^56pN^ beads, but not of phagocytes incubated with PS-TGT^12pN^ beads (Fig. 2, E and F, and movies S6 and S7). In addition, given that constitutively active Rac1 (Rac1(CA)) has been reported to interfere with phagocytic cup closure and inhibit efferocytosis, Rac1(CA) can mitigate the effect of tension of PS-PSRs on engulfment of PS-TGT^56pN^ beads. To test this, we overexpressed Rac1(CA) in phagocytes and evaluated its effect. As expected, engulfment of PS-TGT^56pN^ by phagocytes overexpressing Rac1(CA) was impeded in a manner dependent on the level of Rac1(CA) expression (fig. S2, E and F). These data suggest that tension of PS-PSRs orchestrates phagocytic cup closure via spatiotemporal inactivation of Rac1 during efferocytosis.

Along with Rac1 inactivation, myosin II-mediated contraction is necessary for phagocytic cup closure (*20*). To investigate whether the tension also affects myosin II activity, we performed a frustrated phagocytosis assay that can quantitatively evaluate phagocytic cup extension and contraction (*21*). Macrophages were incubated on PS-TGT-coated plates and their extension and contraction were monitored. During the extension phase, cells spread comparably on PS-TGT^12pN^ and PS-TGT^56pN^ plates. However, during the contraction phase, phagocytes contracted more slowly on the PS-TGT^12pN^ plate than on the PS-TGT^56pN^ plate. In addition, treatment of cells with blebbistatin, a myosin II inhibitor, drastically delayed the contraction of phagocytes on the PS-TGT^56pN^ plate such that it was similar to that on the PS-TGT^12pN^ plate (Fig. 2G and movies S8 to S10). Higher levels of myosin light chain (MLC) phosphorylation were observed around PS-TGT^56pN^ beads and at the leading edge of cells on the PS-TGT^56pN^ plate when macrophages were incubated with PS-TGT beads and on PS-TGT plates, respectively (Fig. 2H, fig. S3, A to C, and movie S11). Overall, these data imply that tension of PS-PSRs causes Rac1 inactivation at the phagocytic cup, leading to both myosin II-mediated contraction and F-actin disassembly required for phagocytic cup closure.

Next, we investigated how tension of PS-PSRs modulates Rac1 activity. Phosphoinositide 3-kinase (PI3K) recruits GTPase-activating proteins (GAPs) for Rac1 by generating PIP3 at the phagocytic cup during efferocytosis, leading to Rac1 inactivation and thus phagocytic cup closure (*22*). Therefore, we reasoned that the tension can alter PI3K activity. To test whether PI3K affects tension tolerance-dependent engulfment, macrophages were treated with LY294002, a PI3K inhibitor, and engulfment of the beads was evaluated. The inhibitor completely abrogated the superior engulfment of PS-TGT^56pN^ beads relative to PS-TGT^12pN^ beads (Fig. 3A), suggesting that PI3K is involved in tension tolerance-dependent engulfment of PS-TGT beads. To further understand the relationship between the tension and PI3K activity, we used the GFP-tagged PH domain of Akt (PH-Akt-GFP) that binds to PIP3, which represents PI3K activity. When PH-Akt-GFP-expressing macrophages were incubated with PS-TGT beads, the GFP signal was higher in phagocytic cups around PS-TGT^56pN^ beads than in phagocytic cups around PS-TGT^12pN^ beads, and this difference was nullified in the presence of the PI3K inhibitor (Fig. 3, B and C, and movies S12 to S14). This implies that tension of PS-PSRs increases the activity of PI3K, leading to Rac1 inactivation in the phagocytic cup.

**Fig. 3.**
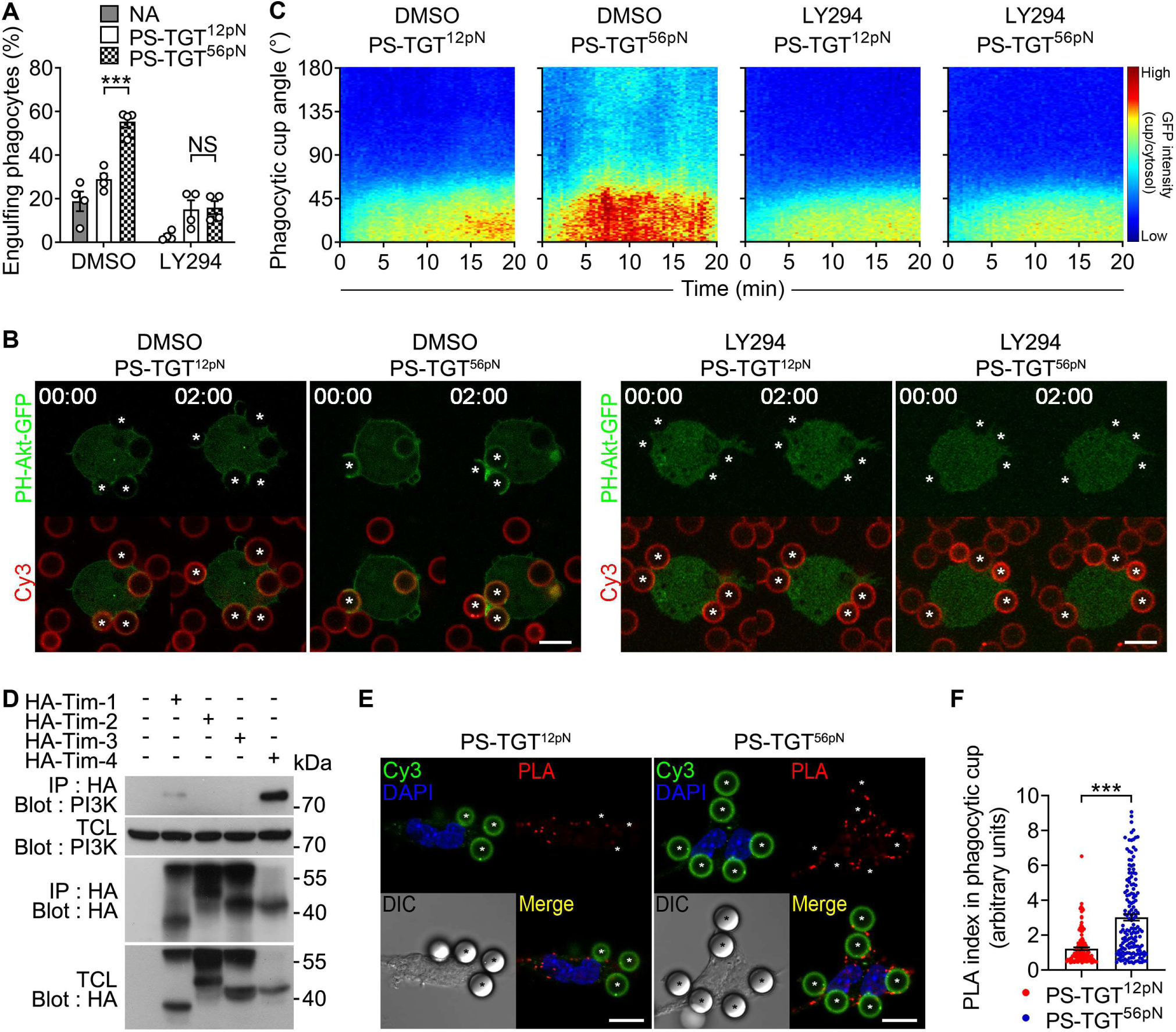
The tension modulates PI3K activity. (**A**) J774A.1 cells pre-incubated with LY294 (LY294002, 50 μM) for 10 min were incubated with the indicated Cy3-labeled beads and analyzed by flow cytometry. Data are mean ± s.e.m. from n=4 independent experiments. ***P < 0.001; NS, not significant; two-way ANOVA. (**B** and **C**) J774A.1 cells stably expressing PH-Akt-GFP were incubated with Cy3-labeled PS-TGT beads for 30 min and observed by time-lapse confocal microscopy. Asterisks indicate beads with extended phagocytic cups (B). The intensities of GFP at phagocytic cups were measured (C). Scale bar, 10 μm. n=67 beads for PS-TGT^12pN^ (DMSO), n=41 beads for PS-TGT^56pN^ (DMSO), n=70 beads for PS-TGT^12pN^ (LY294002), and n=54 beads for PS-TGT^56pN^ (LY294002). (**D**) Lysates of 293T cells transfected with the indicated plasmids were incubated with anti-HA antibody-conjugated agarose beads. Bead-bound proteins were detected by the indicated antibodies. IP, immunoprecipitation; TCL, total cell lysate. (**E** and **F**) Peritoneal macrophages were incubated with Cy3-labeled PS-TGT beads for 10 min, fixed, blocked, and incubated with anti-Tim4 and anti-PI3K antibodies. Then, PLA probes were stained and incubated in amplification solution. PLA signals were observed using confocal microscopy (E), and PLA signals for each bead were quantified (F). Asterisks indicate beads with extended phagocytic cups. Scale bar, 10 μm. Data are mean ± s.e.m. from n=126 beads for PS-TGT^12pN^ and n=153 beads for PS-TGT^56pN^. ***P < 0.001; two-tailed unpaired Student’s *t* test.

We then explored how the tension modulates PI3K activity. Given that PI3K associates with some engulfment receptors (e.g., Tim-1 and Mertk) and all experiments in this study were performed under serum-free conditions in which only direct PSRs could participate in engulfment of the beads (*23–27*), we investigated whether PI3K associates with PSRs and, if so, whether the association is affected by tension of PS-PSRs. PI3K robustly associated with Tim-4 among members of the Tim family and BAI1 (Fig. 3D and fig. 4, A and B). In addition, to confirm the subcellular location of the association between Tim-4 and PI3K at the endogenous level, a proximity ligation assay (PLA) was performed. As a result, in macrophages incubated with PS-TGT^12pN^ beads, PLA signals were not confined to the phagocytic cup and were observed ubiquitously. However, in macrophages incubated with PS-TGT^56pN^ beads, most signals were observed around the phagocytic cup (Fig. 3, E and F). Collectively, these data imply that tension of PS-PSRs increases the association of PI3K with PSRs, e.g., Tim-4, sequentially resulting in Rac1 inactivation, F-actin disassembly, and myosin II-mediated contraction, which ultimately causes phagocytic cup closure.

**Fig. 4.**
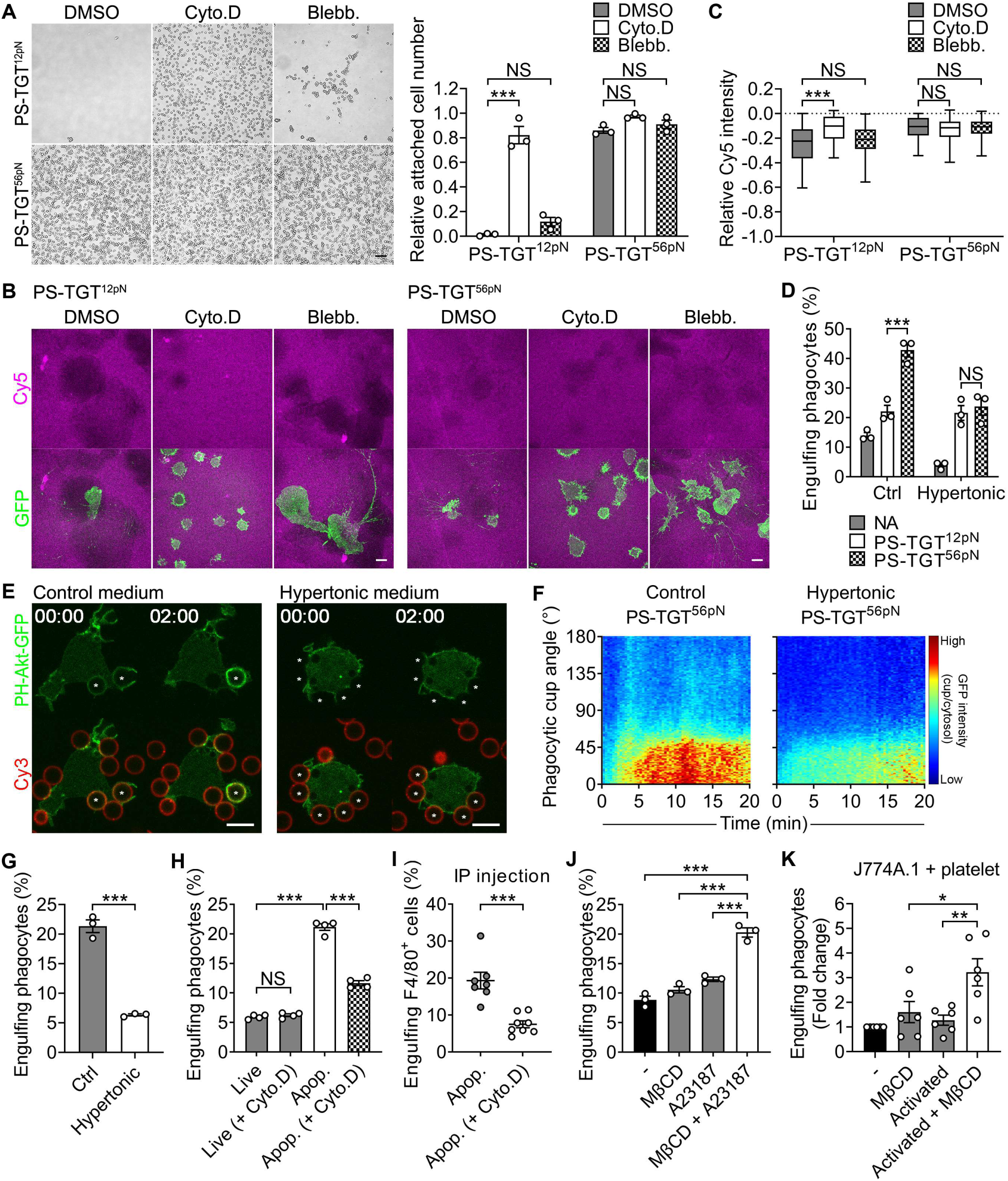
Membrane tension increased by actin polymerization loads tension on PS-PSRs. (**A**) J774A.1 cells incubated on the indicated PS-TGT plates for 5 min were washed and then further incubated in the absence or presence of cytochalasin D (1 μM) or blebbistatin (10 μM) for 60 min. The cell number at 60 min relative to that at 5 min was calculated. Representative images after incubation for 60 min are shown. Scale bar, 200 μm. Cyto.D, cytochalasin D; Blebb, blebbistatin. Data are mean ± s.e.m. from n=3 independent experiments. ***P < 0.001; NS, not significant; two-way ANOVA. (**B** and **C**) J774A.1 cells stably expressing membrane-targeted GFP were incubated on the indicated PS-TGT plates for 60 min and Cy5 fluorescence was observed by confocal microscopy (B). Relative Cy5 intensities were calculated. The intensity of Cy5 in the area where cells were absent was set to 0 (C). Data are presented in the box-and-whisker plot. n=55 cells for PS-TGT^12pN^ (DMSO), n=172 cells for PS-TGT^12pN^ (Cyto.D), n=115 cells for PS-TGT^12pN^ (Blebb.), n=92 cells for PS-TGT^56pN^ (DMSO), n=189 cells for PS-TGT^56pN^ (Cyto.D), and n=161 cells for PS-TGT^56pN^ (Blebb.). ***P < 0.001; NS, not significant; two-way ANOVA. (**D**) J774A.1 cells pre-incubated in medium containing 150 mM sucrose (hypertonic medium) were incubated with the indicated beads for 10 min and analyzed by flow cytometry. Data are mean ± s.e.m. from n=3 independent experiments. ***P < 0.001; NS, not significant; two-way ANOVA. (**E** and **F**) J774A.1 cells stably expressing PH-Akt-GFP were pre-incubated in medium containing 150 mM sucrose and then incubated with Cy3-labeled PS-TGT^56pN^ beads and observed by time-lapse confocal microscopy. Asterisks indicate beads with extended phagocytic cups (E). The intensities of GFP fluorescence at phagocytic cups around the beads were measured (F). Scale bar, 10 μm. n=40 beads for PS-TGT^56pN^ (control medium) and n=51 beads for PS-TGT^56pN^ (hypertonic medium). (**G**) J774A.1 cells pre-incubated in medium containing 150 mM sucrose were incubated with apoptotic Jurkat cells for 30 min and analyzed by flow cytometry. Data are mean ± s.e.m. from n=3 independent experiments. ***P < 0.001; two-tailed unpaired Student’s *t* test. (**H**) J774A.1 cells were incubated with live or apoptotic Jurkat cells treated with or without cytochalasin D (2 μM) for 30 min and analyzed by flow cytometry. Data are mean ± s.e.m. from n=4 independent experiments. ***P < 0.001; NS, not significant; one-way ANOVA. (**I**) TAMRA-stained apoptotic Jurkat cells treated with or without cytochalasin D (2 μM) were intraperitoneally injected into 8-week-old mice. Peritoneal exudates from the mice were stained with an anti-F4/80 antibody and analyzed by flow cytometry. Data are mean ± s.e.m. from n=7 mice for Apop. and n=8 mice for Apop. (+Cyto.D). ***P < 0.001; two-tailed unpaired Student’s *t* test. (**J**) J774A.1 cells were incubated with Jurkat cells treated with MβCD (1 mM), A23187 (10 μM), or MβCD and A23187 for 30 min and analyzed by flow cytometry. Data are mean ± s.e.m. from n=3 independent experiments. ***P < 0.001; one-way ANOVA. (**K**) J774A.1 cells were incubated with platelets treated with MβCD (10 mM) or activated platelets treated with MβCD for 30 min and analyzed by flow cytometry. Data are mean ± s.e.m. from n=6 independent experiments. *P < 0.05, **P < 0.01; one-way ANOVA.

To investigate how tension is loaded on PS-PSRs, actin and myosin were examined because they generate force in cells and are essential for phagocytic cup formation and closure during efferocytosis (*20, 28*). In addition, F-actin disassembly and myosin II-mediated contraction were modulated in a tension tolerance-dependent manner during engulfment of PS-TGT beads (Fig. 2, A, B, G, and H). This suggests that tension of PS-PSRs at the phagocytic cup during efferocytosis can be loaded by actin dynamics and/or myosin II-mediated contraction. To test this possibility, cell detachment from PS-TGT plates was evaluated in the presence or absence of cytochalasin D and blebbistatin, which are inhibitors of actin polymerization and myosin II-mediated contraction, respectively. Cytochalasin D blocked detachment of macrophages from the glass plate to which they were connected via PS-TGT^12pN^ (Fig. 4, A and B) and loss of Cy5 signals in Cy5-labeled PS-TGT^12pN^, but blebbistatin did not (Fig. 4C and fig. S5A). These data imply that actin polymerization, rather than myosin II-mediated contraction, is crucial for tension loading between PS and PSRs at the phagocytic cup. In addition, the tension is not due to an allosteric effect of PSRs induced by PS binding because cytochalasin D still allows PS to bind to its receptors (Fig. 4A).

Given that membrane tension increases during phagocytic cup formation driven by actin polymerization (*29*), this membrane tension may play an important role in loading tension on PS-PSRs. To validate this, we decreased the membrane tension of macrophages using hypertonic solution (*30, 31*). Indeed, the more efficient engulfment of PS-TGT^56pN^ beads relative to PS-TGT^12pN^ beads was abrogated by treating macrophages with hypertonic solution (Fig. 4D). The treatment also reduced the Cy5 MFI of non-engulfing phagocytes upon incubation with PS-TGT^12pN^ beads, but that of non-engulfing phagocytes upon incubation with PS-TGT^56pN^ beads was unaltered (fig. S5B). Furthermore, treatment with hypertonic solution diminished the GFP signal, representing PI3K activity, around PS-TGT^56pN^ beads in macrophages expressing PH-Akt-GFP (Fig. 4, E and F, and movie S15). Consistently, when apoptotic cells were used instead of PS-TGT beads to increase the physiological relevance, phagocytosis of apoptotic cells was drastically impaired when macrophages were treated with hypertonic solution (Fig. 4G). Overall, these data imply that an increase in the membrane tension of phagocytes is coupled with an increase in tension or loading rate exerted on PS-PSRs during efferocytosis.

The membrane stiffness of apoptotic cells is increased (*32, 33*), which is expected to be necessary for loading tension on PS-PSRs during efferocytosis. To apply tension of PS-PSRs, PS on the surface of apoptotic cells must be pulled by PSRs on phagocytes. In this situation, the membrane stiffness of apoptotic cells may be crucial to reach a certain level of tension generated by the pulling of PSRs within the limited phagocytic synaptic space. To validate this, we altered the membrane stiffness of apoptotic cells using cytochalasin D, which decreases membrane stiffness (*33*), and performed an efferocytosis assay. Treatment of apoptotic cells with cytochalasin D substantially impeded efferocytosis *in vitro* and *in vivo* (Fig. 4, H and I).

Notably, the conditioned medium of apoptotic cells treated with cytochalasin D did not affect efferocytosis, demonstrating that the direct effect of cytochalasin D released from apoptotic cells on phagocytes was minimal (fig. S5C). In addition, cytochalasin D did not alter the status of apoptotic cells (fig. S5D). These data imply that increased membrane stiffness in apoptotic cells also contributes to tension generation between PS and PSRs during efferocytosis.

PS is exposed not only on apoptotic cells but also on some live cells, but only apoptotic cells are phagocytosed (*9, 34–36*). In general, this is thought to be due to don’t eat-me signals such as CD47 on live cells (*10*). In addition to these signals, our observations strongly suggest that tension of PS-PSRs is another decisive signal for efferocytosis. To investigate this, we generated PS-exposing Jurkat cells using A23187, a calcium ionophore that stimulates PS exposure on the cell surface without inducing apoptosis (fig. S5, E and F), and artificially increased the membrane stiffness of Jurkat cells by depleting cholesterol using methyl-β-cyclodextrin (MβCD) (*37, 38*). Macrophages efficiently engulfed Jurkat cells treated with both A23187 and MβCD, but engulfment of Jurkat cells treated with A23187 or MβCD alone was minimally increased compared with that of control Jurkat cells (Fig. 4J). To further validate the physiological relevance of these findings, we used platelets, which expose PS on their surface when they are activated for blood coagulation (fig. S5G) (*39, 40*). Engulfment of activated platelets by macrophages was indistinguishable from that of non-activated platelets, but activated platelets treated with MβCD were efficiently engulfed (Fig. 4K). Taken together, these data suggest that tension of PS-PSRs, mediated by both increased membrane tension at the phagocytic cup in phagocytes and membrane stiffness in apoptotic cells, acts as a checkpoint to determine whether a PS-exposing cell should be phagocytosed and this tension is thus essential for completion of efferocytosis.

In sum, we report that efferocytosis, a type of phagocytosis, is a novel example of a process mediated by chemo-mechanical pathways. Upon binding between PS and PSRs, Rac1 is activated, resulting in actin polymerization to form the phagocytic cup around apoptotic cells and thus increasing membrane tension concomitant with tension of PS-PSRs at the phagocytic cup. When the tension reaches a substantial level (e.g., 56 pN), PI3K is recruited to PSRs at the phagocytic cup, which results in Rac1 inactivation, leading to F-actin disassembly and myosin II-mediated contraction required for phagocytic cup closure. Ultimately, tension of PS-PSRs must be generated to successfully complete phagocytosis of apoptotic cells (fig. S6A).

Our findings provide insights into how chemical and mechanical signals generated during phagocytosis of PS-exposing cells are delicately coordinated to enhance the integrity of efferocytosis without errors. More specifically, biochemical signaling of efferocytosis initiated by binding of PS to PSRs is converted into a mechanical signal (i.e., tension of PS-PSRs). The tension is eventually converted back into a biochemical signal that determines whether PS-exposing cells being engulfed are completely phagocytosed (fig. S6B). By generating a mechanical signal during phagocytosis of PS-exposing cells, phagocytes can avoid removing all PS-exposing cells and precisely remove cells to be cleared, e.g., apoptotic cells. Indeed, many live cells expose PS on their surfaces including some cancer cells (*34–36, 41–44*). Cancer immunotherapy based on innate immunity has recently attracted much attention (*45*). Cancer cells may evade phagocytosis by immune cells via nullifying this type of mechanical signal as a checkpoint during phagocytosis. It would be interesting to investigate the correlation of cancer cell phagocytosis with tension between a ligand on cancer cells and its corresponding receptor on immune cells, which may provide insight to develop cancer immunotherapy.

## Acknowledgments

We thank Michiyuki Matsuda (Kyoto University) and Jihye Seong (Korea Institute of Science and Technology) for kindly providing the plasmids of Raichu-Rac1 and PH-Akt-GFP, respectively.

## Author contributions

Conceptualization, G.L. and D.P.; Methodology, C.M., H.C, J.Y., B.M., J.J., N.S., H.M., S.Y., J.L., and M.H.; Formal analysis, C.M., H.C, J.Y., P.J., G.L., and D.P.; Resources, J.J. and W.-J.C.; Writing, C.M., H.C, J.Y., G.L., and D.P.; Funding acquisition, G.L. and D.P.

## Data and materials availability

Data from this study are available within the main text and the supplementary materials or from the corresponding authors (G.L. and D.P.) upon reasonable request.

## Supplementary Materials

Materials and Methods

Supplementary Text (Uncropped blots of western blot panels, Gating strategies used for flow cytometry analysis, Movie legends, Synthesis of 18:1-12:0(maleimide) phosphatidylcholine)

Figs. S1 to S6

Movies S1 to S15

## References and Notes

1. S. Arandjelovic, K. S. Ravichandran, Phagocytosis of apoptotic cells in homeostasis. Nat Immunol 16, 907–917 (2015).

2. A. C. Doran, A. Yurdagul, Jr., I. Tabas, Efferocytosis in health and disease. Nat Rev Immunol 20, 254–267 (2020).

3. V. A. Fadok, D. L. Bratton, S. C. Frasch, M. L. Warner, P. M. Henson, The role of phosphatidylserine in recognition of apoptotic cells by phagocytes. Cell Death Differ 5, 551–562 (1998).

4. V. A. Fadok et al., Exposure of phosphatidylserine on the surface of apoptotic lymphocytes triggers specific recognition and removal by macrophages. J Immunol 148, 2207–2216 (1992).

5. P. A. Oldenborg et al., Role of CD47 as a marker of self on red blood cells. Science 288, 2051–2054 (2000).

6. A. A. Barkal et al., CD24 signalling through macrophage Siglec-10 is a target for cancer immunotherapy. Nature 572, 392–396 (2019).

7. S. Brown et al., Apoptosis disables CD31-mediated cell detachment from phagocytes promoting binding and engulfment. Nature 418, 200–203 (2002).

8. A. A. Barkal et al., Engagement of MHC class I by the inhibitory receptor LILRB1 suppresses macrophages and is a target of cancer immunotherapy. Nat Immunol 19, 76–84 (2018).

9. K. Segawa, J. Suzuki, S. Nagata, Constitutive exposure of phosphatidylserine on viable cells. Proc Natl Acad Sci U S A 108, 19246–19251 (2011).

10. S. Jaiswal et al., CD47 is upregulated on circulating hematopoietic stem cells and leukemia cells to avoid phagocytosis. Cell 138, 271–285 (2009).

11. J. M. Kinchen, K. S. Ravichandran, Journey to the grave: signaling events regulating removal of apoptotic cells. J Cell Sci 120, 2143–2149 (2007).

12. R. K. Tsai, D. E. Discher, Inhibition of “self” engulfment through deactivation of myosin-II at the phagocytic synapse between human cells. J Cell Biol 180, 989–1003 (2008).

13. S. Mylvaganam, S. A. Freeman, S. Grinstein, The cytoskeleton in phagocytosis and macropinocytosis. Curr Biol 31, R619–R632 (2021).

14. X. Wang, T. Ha, Defining single molecular forces required to activate integrin and notch signaling. Science 340, 991–994 (2013).

15. R. Roy, S. Hohng, T. Ha, A practical guide to single-molecule FRET. Nat Methods 5, 507–516 (2008).

16. M. Nakaya, M. Kitano, M. Matsuda, S. Nagata, Spatiotemporal activation of Rac1 for engulfment of apoptotic cells. Proc Natl Acad Sci U S A 105, 9198–9203 (2008).

17. H. Moon et al., Crbn modulates calcium influx by regulating Orai1 during efferocytosis. Nat Commun 11, 5489 (2020).

18. M. R. Elliott, K. M. Koster, P. S. Murphy, Efferocytosis Signaling in the Regulation of Macrophage Inflammatory Responses. J Immunol 198, 1387–1394 (2017).

19. R. E. Itoh et al., Activation of rac and cdc42 video imaged by fluorescent resonance energy transfer-based single-molecule probes in the membrane of living cells. Mol Cell Biol 22, 6582–6591 (2002).

20. Y. Ikeda et al., Rac1 switching at the right time and location is essential for Fcgamma receptor-mediated phagosome formation. J Cell Sci 130, 2530–2540 (2017).

21. D. T. Kovari et al., Frustrated Phagocytic Spreading of J774A-1 Macrophages Ends in Myosin II-Dependent Contraction. Biophys J 111, 2698–2710 (2016).

22. D. Schlam et al., Phosphoinositide 3-kinase enables phagocytosis of large particles by terminating actin assembly through Rac/Cdc42 GTPase-activating proteins. Nat Commun 6, 8623 (2015).

23. P. Sen et al., Apoptotic cells induce Mer tyrosine kinase-dependent blockade of NF-kappaB activation in dendritic cells. Blood 109, 653–660 (2007).

24. A. J. de Souza et al., T cell Ig and mucin domain-1-mediated T cell activation requires recruitment and activation of phosphoinositide 3-kinase. J Immunol 180, 6518–6526 (2008).

25. G. X. Ruan, A. Kazlauskas, Axl is essential for VEGF-A-dependent activation of PI3K/Akt. EMBO J 31, 1692–1703 (2012).

26. M. Miyanishi et al., Identification of Tim4 as a phosphatidylserine receptor. Nature 450, 435–439 (2007).

27. D. Park et al., BAI1 is an engulfment receptor for apoptotic cells upstream of the ELMO/Dock180/Rac module. Nature 450, 430–434 (2007).

28. K. Carvalho et al., Actin polymerization or myosin contraction: two ways to build up cortical tension for symmetry breaking. Philos Trans R Soc Lond B Biol Sci 368, 20130005 (2013).

29. V. Jaumouille, C. M. Waterman, Physical Constraints and Forces Involved in Phagocytosis. Front Immunol 11, 1097 (2020).

30. A. R. Houk et al., Membrane tension maintains cell polarity by confining signals to the leading edge during neutrophil migration. Cell 148, 175–188 (2012).

31. T. A. Masters, B. Pontes, V. Viasnoff, Y. Li, N. C. Gauthier, Plasma membrane tension orchestrates membrane trafficking, cytoskeletal remodeling, and biochemical signaling during phagocytosis. Proc Natl Acad Sci U S A 110, 11875–11880 (2013).

32. N. I. Nikolaev, T. Muller, D. J. Williams, Y. Liu, Changes in the stiffness of human mesenchymal stem cells with the progress of cell death as measured by atomic force microscopy. J Biomech 47, 625–630 (2014).

33. W. A. Lam, M. J. Rosenbluth, D. A. Fletcher, Chemotherapy exposure increases leukemia cell stiffness. Blood 109, 3505–3508 (2007).

34. J. I. Elliott et al., Membrane phosphatidylserine distribution as a non-apoptotic signalling mechanism in lymphocytes. Nat Cell Biol 7, 808–816 (2005).

35. S. Riedl et al., In search of a novel target - phosphatidylserine exposed by non-apoptotic tumor cells and metastases of malignancies with poor treatment efficacy. Biochim Biophys Acta 1808, 2638–2645 (2011).

36. S. M. Qadri, R. Bissinger, Z. Solh, P. A. Oldenborg, Eryptosis in health and disease: A paradigm shift towards understanding the (patho)physiological implications of programmed cell death of erythrocytes. Blood Rev 31, 349–361 (2017).

37. J. Suzuki, M. Umeda, P. J. Sims, S. Nagata, Calcium-dependent phospholipid scrambling by TMEM16F. Nature 468, 834–838 (2010).

38. A. Biswas, P. Kashyap, S. Datta, T. Sengupta, B. Sinha, Cholesterol Depletion by MbetaCD Enhances Cell Membrane Tension and Its Variations-Reducing Integrity. Biophys J 116, 1456–1468 (2019).

39. S. M. Schoenwaelder et al., Two distinct pathways regulate platelet phosphatidylserine exposure and procoagulant function. Blood 114, 663–666 (2009).

40. B. R. Lentz, Exposure of platelet membrane phosphatidylserine regulates blood coagulation. Prog Lipid Res 42, 423–438 (2003).

41. K. Fischer et al., Antigen recognition induces phosphatidylserine exposure on the cell surface of human CD8+ T cells. Blood 108, 4094–4101 (2006).

42. J. I. Elliott et al., Phosphatidylserine exposure in B lymphocytes: a role for lipid packing. Blood 108, 1611–1617 (2006).

43. K. J. de Vries, T. Wiedmer, P. J. Sims, B. M. Gadella, Caspase-independent exposure of aminophospholipids and tyrosine phosphorylation in bicarbonate responsive human sperm cells. Biol Reprod 68, 2122–2134 (2003).

44. J. H. Stafford, P. E. Thorpe, Increased exposure of phosphatidylethanolamine on the surface of tumor vascular endothelium. Neoplasia 13, 299–308 (2011).

45. M. Feng et al., Phagocytosis checkpoints as new targets for cancer immunotherapy. Nat Rev Cancer 19, 568–586 (2019).

