## Supplementary materials for "Molecular tension authenticates apoptotic cells being phagocytosed"

### Figure S1

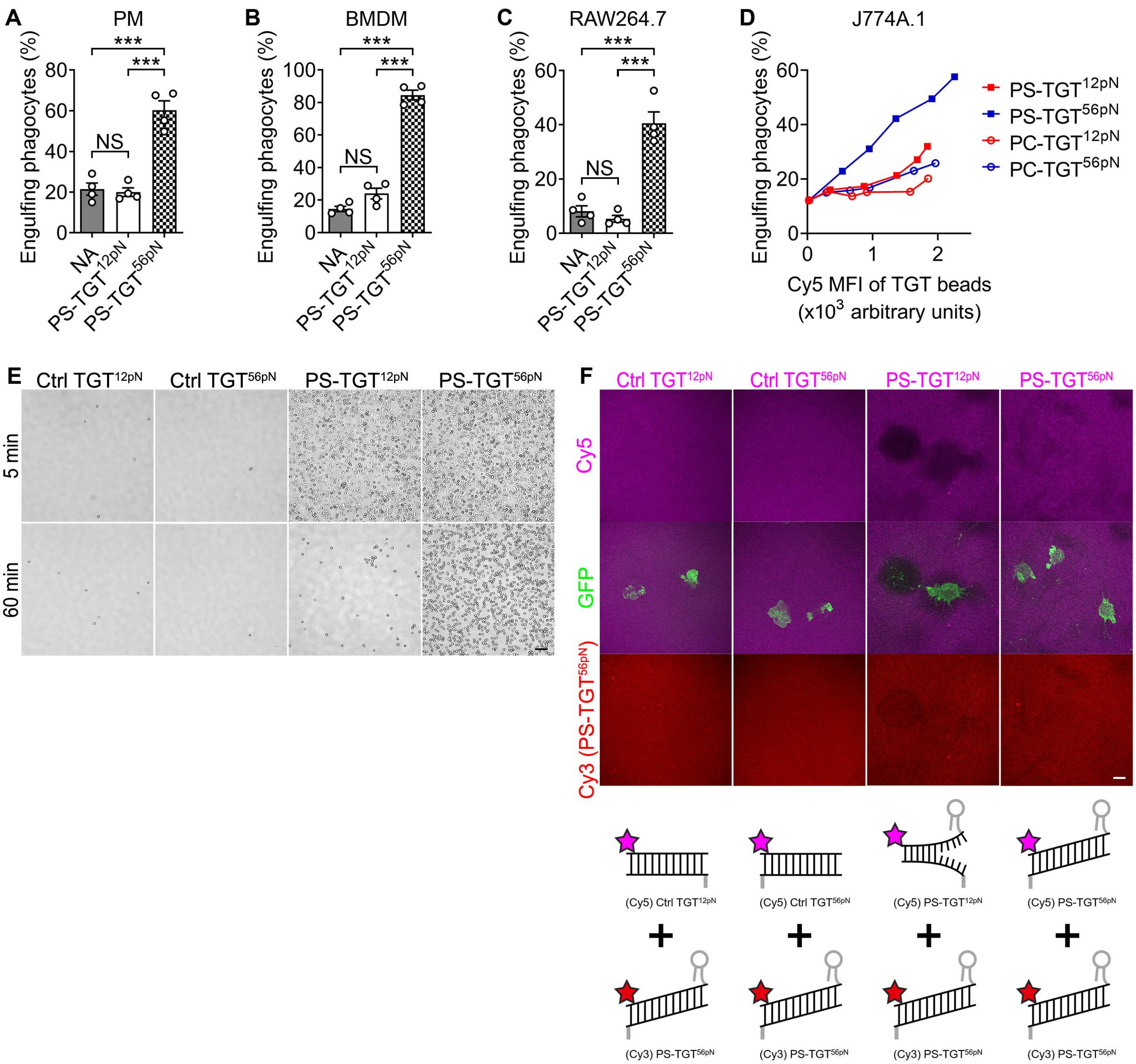

A
PM
Engulfing phagocytes (%)
NS
***
NA
PS-TGT12pN
PS-TGT56pN
B
BMDM
C
RAW264.7
D
J774A.1
Cy5 MFI of TGT beads
(x10^3 arbitrary units)
PS-TGT12pN
PS-TGT56pN
PC-TGT12pN
PC-TGT56pN
E
Ctrl TGT12pN
Ctrl TGT56pN
5 min
60 min
F
Cy5
GFP
Cy3 (PS-TGT56pN)
(Cy5) Ctrl TGT12pN
(Cy5) Ctrl TGT56pN
(Cy5) PS-TGT12pN
(Cy5) PS-TGT56pN
(Cy3) PS-TGT56pN

**Fig. S1. Tension-dependent engulfment of PS-TGT beads by various macrophages**

(**A** to **C**) Peritoneal macrophages (A), bone marrow-derived macrophages (B), or RAW264.7 cells (C) were incubated with the indicated beads for 10 min and analyzed by flow cytometry. NA, neutravidin bead; PM, peritoneal macrophages; BMDM, bone marrow-derived macrophages. Data are mean ± s.e.m. from n=4 independent experiments. ***P < 0.001; NS, not significant; one-way ANOVA. (**D**) Various concentrations of the indicated TGT were linked to neutravidin beads. J774A.1 cells were incubated with the beads for 10 min and analyzed by flow cytometry. (**E**) The extended figure for Fig. 1F. J774A.1 cells incubated on PS-TGT plates for 5 min were washed, further incubated for 60 min, and washed. Scale bar, 200 μm. (**F**) The extended figure for Fig. 1G. J774A.1 cells stably expressing membrane-targeted GFP were incubated on the indicated glass plates, on which a mixture of the indicated Cy5-labeled TGTs and Cy3-labeled PS-TGT$^{56pN}$ was immobilized, for 60 min and observed by confocal microscopy (top). Schematic diagram of a mixture of the indicated Cy5-labeled TGTs and Cy3-labeled PS-TGT$^{56pN}$ immobilized on glass plates (bottom). Scale bar, 10 μm.

Figure S2

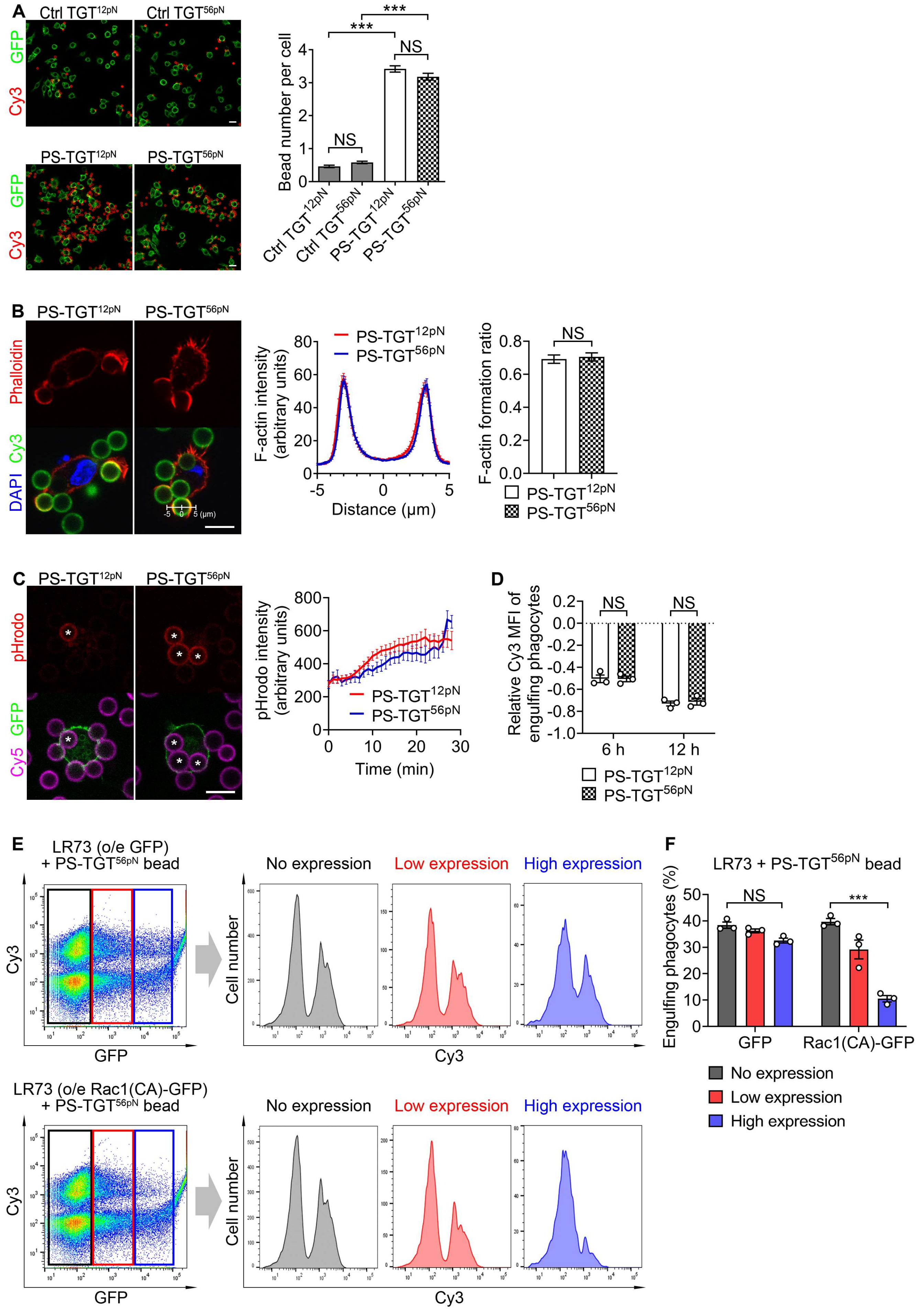
A
Ctrl TGT12pN
Ctrl TGT56pN
PS-TGT12pN
PS-TGT56pN
GFP
Cy3
Bead number per cell
NS
***
B
Phalloidin
Cy3
DAPI
F-actin intensity (arbitrary units)
Distance (μm)
F-actin formation ratio
C
pHrodo
GFP
Cy5
pHrodo intensity (arbitrary units)
Time (min)
D
Relative Cy3 MFI of engulfing phagocytes
6 h
12 h
E
LR73 (o/e GFP) + PS-TGT56pN bead
LR73 (o/e Rac1(CA)-GFP) + PS-TGT56pN bead
No expression
Low expression
High expression
Cell number
F
LR73 + PS-TGT56pN bead
Engulfing phagocytes (%)
GFP
Rac1(CA)-GFP

**Fig. S2. The tension does not affect degradation or binding of PS-TGT beads.**

(**A**) J774A.1 cells were incubated with the indicated beads at 4°C for 30 min and the number of bound beads per phagocyte was quantified. Scale bar, 20 μm. Data are mean ± s.e.m. from n=512 cells for Ctrl $TGT^{12pN}$, n=616 cells for Ctrl $TGT^{56pN}$, n=506 cells for PS-$TGT^{12pN}$, and n=443 cells for PS-$TGT^{56pN}$. ***$P < 0.001$; NS, not significant; one-way ANOVA. (**B**) J774A.1 cells were incubated with Cy3-labeled PS-TGT beads for 10 min, stained with Alexa Fluor 594-conjugated phalloidin, and observed by confocal microscopy (left). Scale bar, 10 μm. The intensities of F-actin across the phagocytic cup were analyzed by a line scan of the phagocytic cup around the beads. Data are mean ± s.e.m. from n=45 beads for PS-$TGT^{12pN}$ and n=40 beads for PS-$TGT^{56pN}$ (middle). The F-actin formation ratio was calculated as the ratio of the number of beads with F-actin to the total number of beads attached to the cell. Data are mean ± s.e.m. from n=121 cells for PS-$TGT^{12pN}$ and n=123 cells for PS-$TGT^{56pN}$. NS, not significant; two-tailed unpaired Student's *t* test (right). (**C**) J774A.1 cells stably expressing membrane-targeted GFP were incubated with pHrodo-labeled and Cy5-labeled PS-TGT beads and observed by time-lapse confocal microscopy for 30 min. Asterisks indicate completely ingested beads. The intensities of pHrodo were measured after cells had completely engulfed the beads (right). Scale bar, 10 μm. Data are mean ± s.e.m. from n=41 beads for PS-$TGT^{12pN}$ and n=24 beads for PS-$TGT^{56pN}$. (**D**) J774A.1 cells incubated with Cy3-labeled PS-TGT beads for 10 min were washed, further incubated for the indicated durations, and analyzed by flow cytometry. The Cy3 MFI of engulfing phagocytes at 10 min was set to 0. Data are mean ± s.e.m. from n=3 independent experiments. NS, not significant; two-way ANOVA. (**E** and **F**) LR73 cells transfected with GFP or Rac1(CA)-GFP were incubated for 2 h with Cy3-labeled PS-$TGT^{56pN}$ beads and analyzed by flow cytometry. The color boxes in the dot plots indicate GFP expression levels (E). Data are mean ± s.e.m. from n=3 independent experiments. ***$P < 0.001$; NS, not significant; two-way ANOVA.

### Figure S3

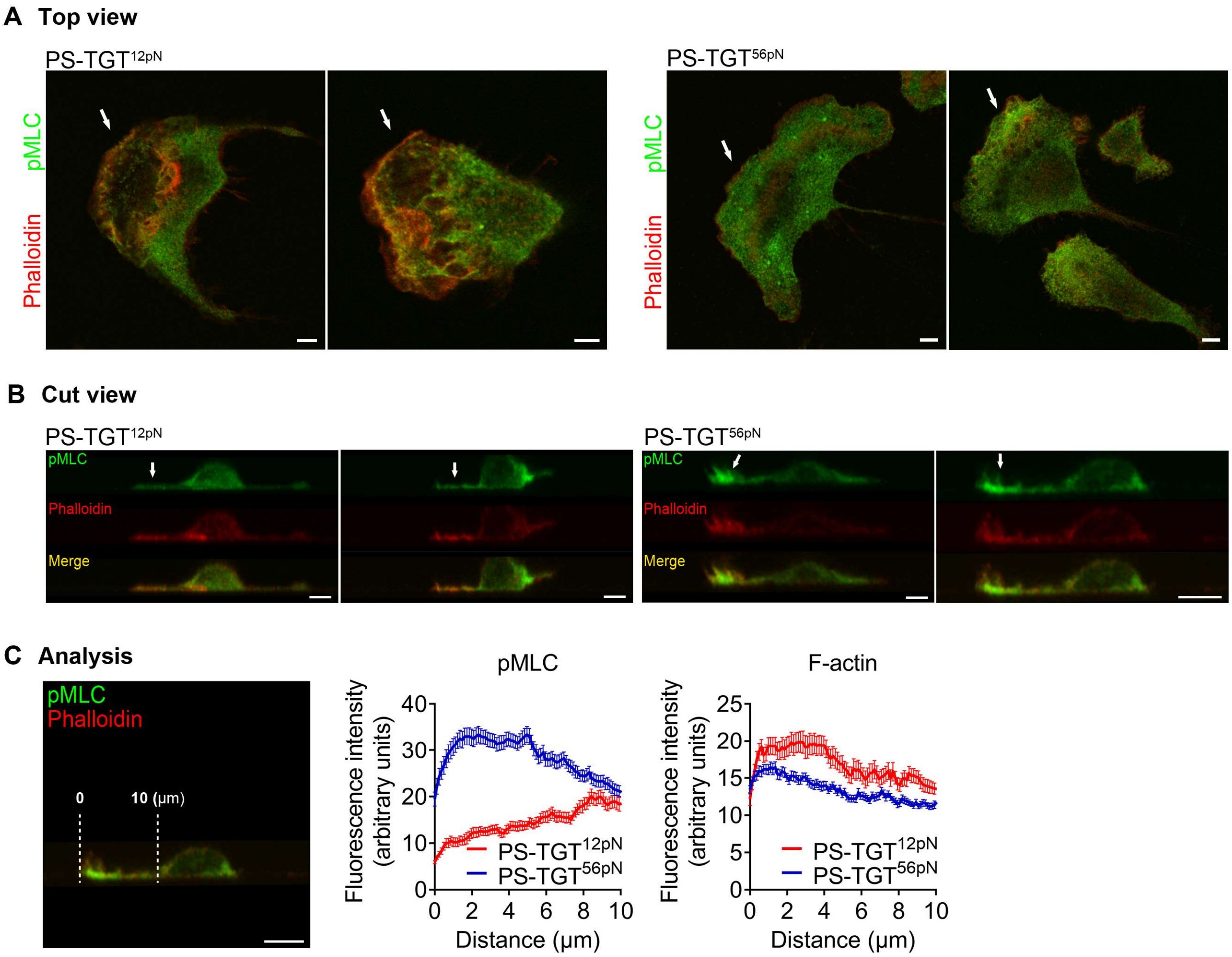

A Top view
PS-TGT12pN
PS-TGT56pN
pMLC
Phalloidin
B Cut view
PS-TGT12pN
PS-TGT56pN
pMLC
Phalloidin
Merge
C Analysis
pMLC
Phalloidin
0
10 (μm)
pMLC
F-actin
Fluorescence intensity (arbitrary units)
Distance (μm)
PS-TGT12pN
PS-TGT56pN

**Fig. S3. PS-TGT with high tension tolerance increases the level of phospho-MLC.**

(**A** to **C**) J774A.1 cells were incubated on glass plates, on which the indicated PS-TGTs were immobilized, washed, stained with an anti-pMLC antibody and phalloidin, and observed by confocal microscopy. The intensities of F-actin and pMLC staining were measured. Arrows indicate actin-enriched regions (the leading edge of cells). Scale bar, 5 μm. Data are mean ± s.e.m. from n=46 cells for PS-$TGT^{12pN}$ and n=67 cells for PS-$TGT^{56pN}$.

### Figure S4

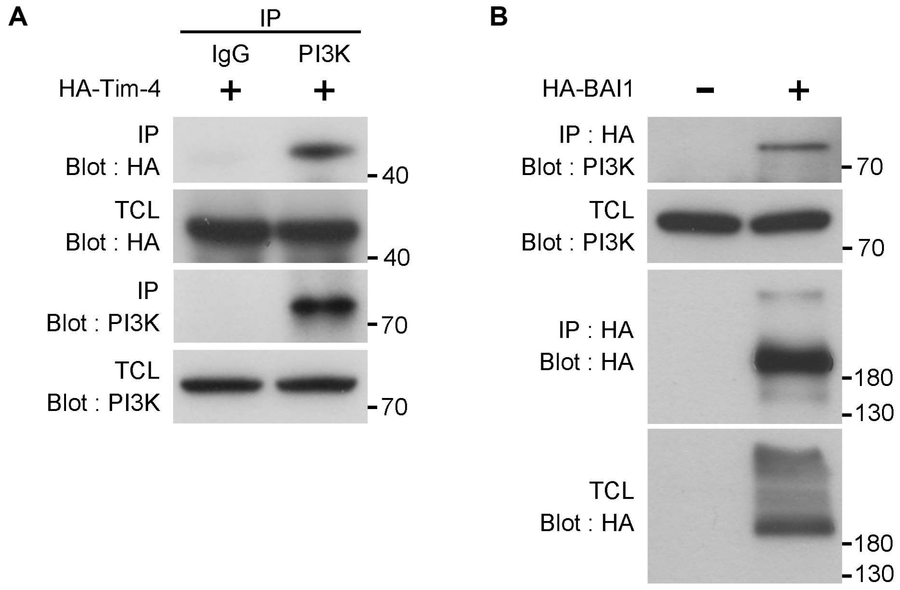
A
IP
IgG
PI3K
HA-Tim-4
+
+
IP
Blot : HA
40
TCL
Blot : HA
40
IP
Blot : PI3K
70
TCL
Blot : PI3K
70
B
HA-BAI1
-
+
IP : HA
Blot : PI3K
70
TCL
Blot : PI3K
70
IP : HA
Blot : HA
180
130
TCL
Blot : HA
180
130

**Fig. S4. PI3K associates with PSRs.**

(**A** and **B**) 293T cells were transfected with the indicated plasmids and lysed. PI3K or HA-BAI1 was precipitated with an anti-PI3K antibody (A) or an anti-HA antibody (B), respectively. Proteins precipitated by the antibody were detected by the indicated antibodies. IP, immunoprecipitation; TCL, total cell lysate.

Figure S5

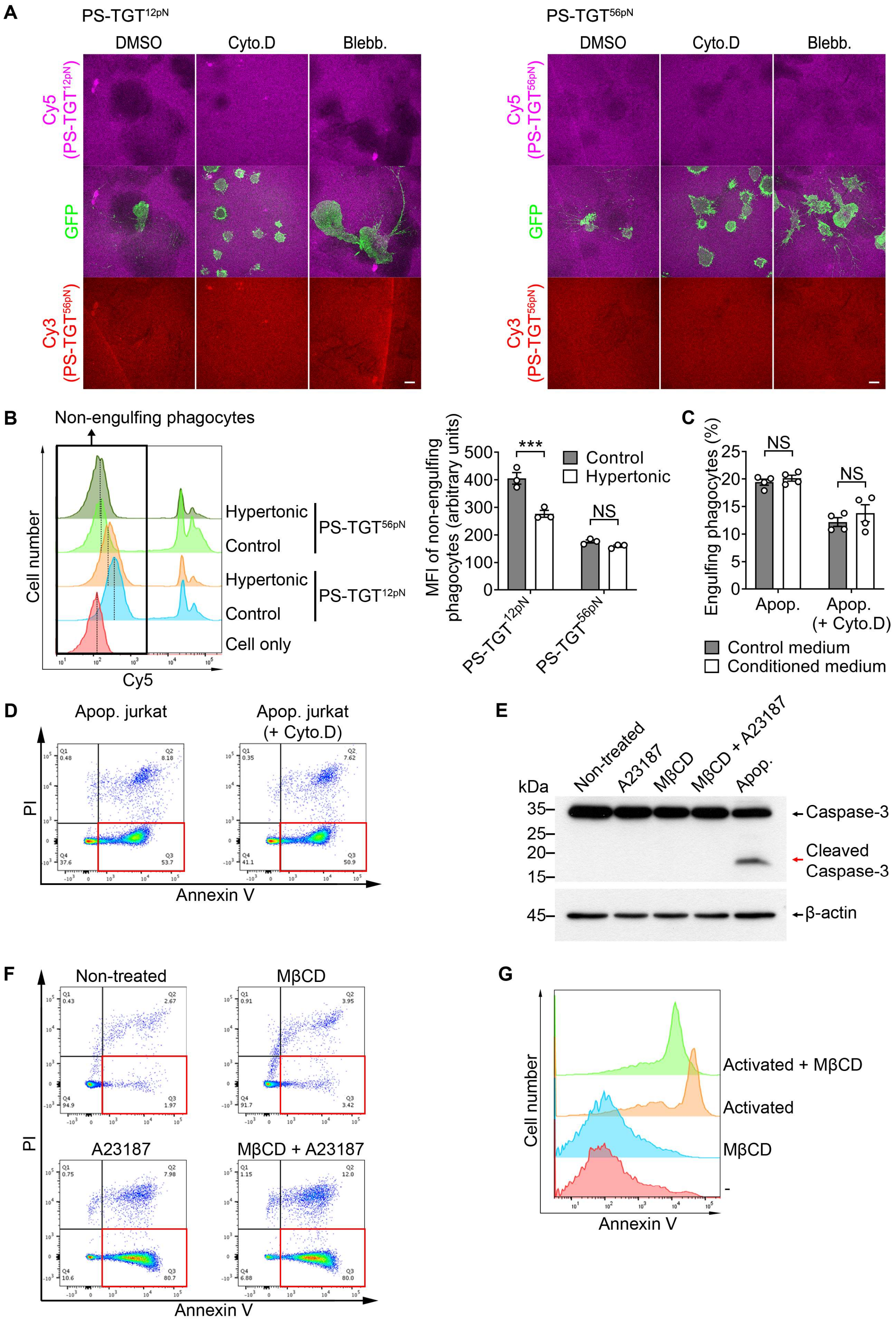

A
PS-TGT12pN
DMSO
Cyto.D
Blebb.
PS-TGT56pN
Cy5 (PS-TGT12pN)
GFP
Cy3 (PS-TGT56pN)
Cy5 (PS-TGT56pN)
B
Non-engulfing phagocytes
Cell number
Hypertonic
Control
PS-TGT56pN
PS-TGT12pN
Cell only
Cy5
MFI of non-engulfing phagocytes (arbitrary units)
***
NS
Control
Hypertonic
C
Engulfing phagocytes (%)
Apop.
Apop. (+ Cyto.D)
Control medium
Conditioned medium
D
Apop. jurkat
Apop. jurkat (+ Cyto.D)
PI
Annexin V
E
kDa
Non-treated
A23187
MβCD
MβCD + A23187
Apop.
Caspase-3
Cleaved Caspase-3
β-actin
F
Non-treated
MβCD
A23187
MβCD + A23187
G
Activated + MβCD
Activated
MβCD
Cell number
Annexin V

**Fig. S5. Actin-mediated membrane tension generates tension between PS and its receptors.**

(**A**) The extended figure for Fig. 4B. J774A.1 cells stably expressing membrane-targeted GFP were incubated on glass plates coated with mixtures of Cy3-labeled PS-TGT$^{56pN}$ and the indicated Cy5-labeled PS-TGTs for 60 min and observed by confocal microscopy. Cyto.D, cytochalasin D; Blebb, blebbistatin. (**B**) J774A.1 cells pre-incubated in control medium or medium containing 150 mM sucrose (hypertonic medium) were incubated with the indicated beads for 10 min and analyzed by flow cytometry. Representative histograms of phagocytes (left) and MFIs of non-engulfing phagocytes (right) are shown. The box and dotted lines in the histograms indicate non-engulfing phagocytes and the peak of the fluorescence intensity of non-engulfing phagocytes, respectively. Data are mean ± s.e.m. from n=3 independent experiments. ***$P < 0.001$; NS, not significant; two-way ANOVA. (**C**) Apoptotic Jurkat cells treated with or without cytochalasin D (2 μM) were incubated on a clean bench for 20 min and the supernatant was collected as conditioned medium after centrifugation. Then, apoptotic Jurkat cells were resuspended in control medium or the conditioned medium and fed to J774A.1 cells for 30 min. Engulfing phagocytes were analyzed by flow cytometry. Data are mean ± s.e.m. from n=4 independent experiments. NS, not significant; two-way ANOVA. (**D**) Apoptotic Jurkat cells treated with or without cytochalasin D were stained with PI and Annexin V and analyzed by flow cytometry. (**E**) Apoptotic Jurkat cells and Jurkat cells treated with A23187, MβCD, or A23187 and MβCD were lysed. Caspase-3 in the lysates was detected by immunoblotting using an anti-Caspase-3 antibody. (**F**) Jurkat cells treated with the indicated drugs were stained with PI and Annexin V and analyzed by flow cytometry. (**G**) Platelets treated with MβCD (10 mM) or activated platelets treated with MβCD for 30 min were stained with Annexin V and analyzed by flow cytometry.

Figure S6

A

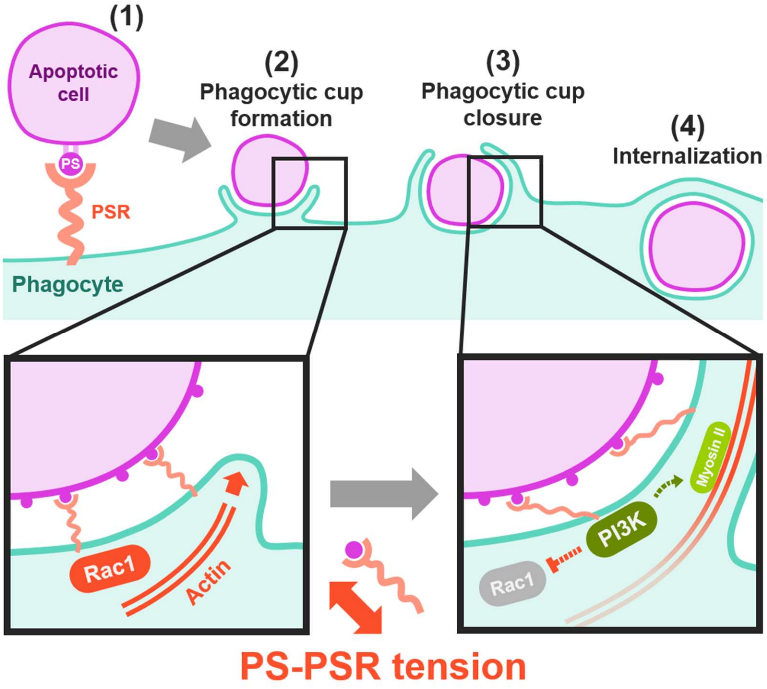


B

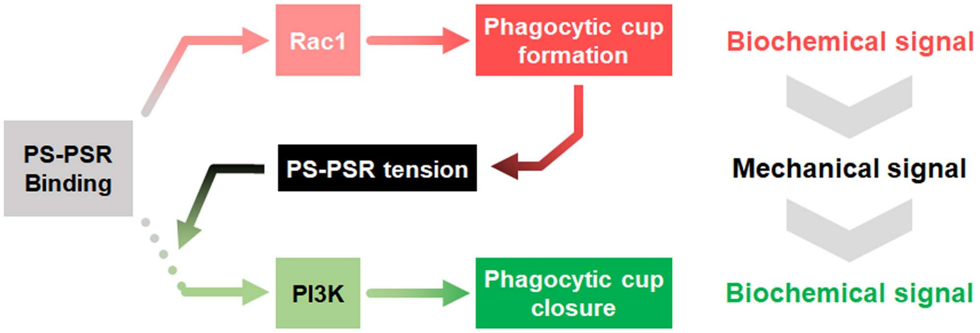


C 18:1-12:0(maleimide) PS

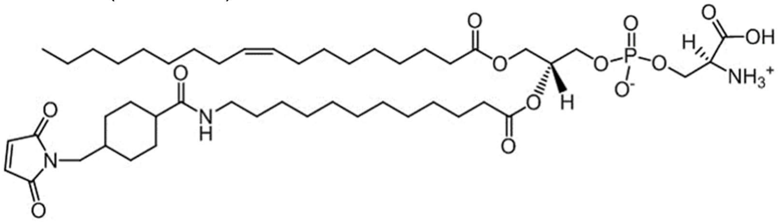

D 18:1-12:0(maleimide) PC

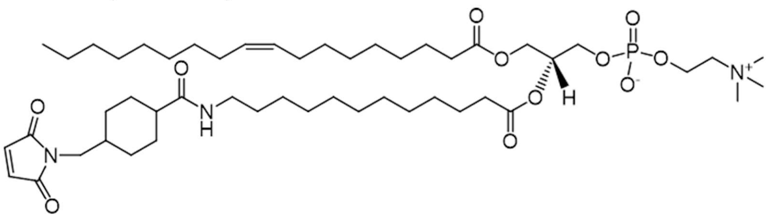

**Fig. S6. A chemo-mechanical signaling pathway in efferocytosis and structures of PS- and PC-maleimide**

(**A**) Schematic diagram of a chemo-mechanical signaling pathway in efferocytosis. Upon binding of PS to PSRs (1), activated Rac1 induces actin polymerization to form the phagocytic cup, which results in an increase of membrane tension concomitant with tension of PS-PSRs in the engulfment synapse (2). Then, the tension causes PI3K to be recruited to PSRs, leading to Rac1 inactivation, which induces F-actin disassembly and myosin II-mediated contraction required for phagocytic cup closure (3). These ultimately complete ingestion of apoptotic cells by phagocytes (4). (**B**) The interconversion between chemical and mechanical signals during efferocytosis. Efferocytosis is initiated by the binding of PS to PSRs, which activates Rac1. Rac1 activation induces actin polymerization, leading to phagocytic cup formation and an increase in membrane tension, which generates tension on PS-PSRs along with the increased stiffness of the apoptotic cell membrane. As the tension escalates to significant levels (e.g., 56 pN), it initiates the recruitment of PI3K to PSRs at the phagocytic cup. This recruitment results in Rac1 inactivation, facilitating the disassembly of F-actin and the myosin II-driven contraction required for phagocytic cup closure. (**C** and **D**) Structures of PS and PC covalently linked with maleimide. Maleimide was linked to the lauric acid of PS (C) or PC (D).

#### Materials and Methods

##### Plasmids and reagents

Raichu-Rac1 (Raichu-Rac1 1011x) and PH-Akt-GFP (Addgene #51465) were generous gifts from Michiyuki Matsuda (Kyoto University) and Jihye Seong (Korea Institute of Science and Technology), respectively. The Cys-Val-Ile-Met sequence was introduced after GFP in the pEGFP-N1 vector to generate GFP-CAAX and sequenced to confirm the fidelity of the introduced sequence. The antibodies used in the study were anti-Tim-4 (NBP1-76702, Novus Biologicals), anti-PI3K (ab86714, Abcam), anti-HA (#3724, Cell Signaling Technology), anti-Caspase-3 (#9622, Cell Signaling Technology), Alexa Fluor 488-conjugated goat anti-mouse (A11029, Invitrogen), Alexa Fluor 647-conjugated donkey anti-rabbit (A31573, Invitrogen), anti-MLC (ab2480, Abcam), Alexa Fluor 488-conjugated goat anti-rabbit (A11008, Invitrogen), and anti-mouse F4/80 (123108, BioLegend). LY294002 (70920, Cayman Chemical Company), phalloidin (A12381, Invitrogen), cytochalasin D (C8273, Sigma-Aldrich), blebbistatin (B0560, Sigma-Aldrich), A23187 (C7522, Sigma-Aldrich), MβCD (C4555, Sigma-Aldrich), citrate-dextrose (ACD; C3821, Sigma-Aldrich), prostaglandin E1 (PGE1; P5515, Sigma-Aldrich), thrombin (T7326, Sigma-Aldrich), and collagen (631-00771, Nitta Gelatin) were purchased.

##### Cell culture and transfection

J774A.1 (DMEM), RAW264.7 (DMEM), Jurkat (RPMI), and 293T (DMEM) cells were maintained in the specified medium containing 10% fetal bovine serum and 1% penicillin-streptomycin-glutamine. J774A.1 cells were transfected with FuGENE® HD transfection reagent (E2311, Promega). To generate J774A.1 cells stably expressing Raichu-Rac1, PH-Akt-GFP, or membrane-targeted GFP (GFP-CAAX), J774A.1 cells were transfected with pApuro and the indicated plasmids and maintained in medium containing puromycin. The stable expression and subcellular localization of the proteins in the cells were confirmed by flow cytometry and confocal microscopy.

**Mice**

C57BL/C were purchased from Taconic bioscience. The mice were bred in equipped animal facility with temperature at 20–25 °C and humidity at 30–70%, under the same dark/light cycle (12:12). All experiments using mice in the study were approved by the animal care and ethics committees of GIST in accordance with the national institutes of health guide for the care and use of laboratory animals.

**Synthesis of PS-TGT and PC-TGT**

TGT synthesis was previously reported (*14*). Briefly, thiol-modified single-stranded DNA (ssDNA) was purchased from Integrated DNA Technologies (upper strand, 5'-/5AmMC6/CAC AGC ACG GAG GCA CGA CAC/3ThioMC3-D/-3'; lower strand, 12 pN: 5'-/5Biosg/GTG TCG TGC CTC CGT GCT GTG-3' and 56 pN: 5'-GTG TCG TGC CTC CGT GCT GTG/3Bio/-3'). The upper strand was labeled with NHS-ester Cy3 (PA13101, Cytiva) or Cy5 (PA15101, Cytiva) via 5AmMC6 modification of the upper strand. Dye labeling buffer contained 100 mM sodium tetraborate decahydrate with DEPC (pH 8.5). The ssDNA (5 nmol) and NHS-ester Cy3 or Cy5 (0.1 mg) were mixed together and incubated overnight at 4°C with vortexing. Then, dye-labeled ssDNA was precipitated using ethanol. The thiol group of ssDNA must be deprotected by 0.5 μM TCEP and degassed for 20 min before thiol-maleimide conjugation. Customized 18:1-12:0(maleimide) PS (Avanti Polar Lipids) or synthesized 18:1-12:0(maleimide) PC (fig. S6, C and D) with DMSO was mixed with the upper strand at a molar ratio of 1:40 and incubated overnight at 25°C. Then, PS- or PC-conjugated ssDNA was separated by electrophoresis in a 20% native gel, purified by ethanol precipitation, and annealed with the lower strands.

**Preparation of TGT beads and plates**

To prepare 2×10$^6$ TGT beads, 50 μL neutravidin-coated polystyrene particles (6.0–8.0 μm; NVP-60-5, Spherotech) were washed and resuspended in 50 μL DMEM containing 5 mg/mL bovine serum albumin (BSA). The beads were then incubated with 5 μL of 5 μM TGT at 4°C for 30 min, washed, and resuspended in DMEM containing 5 mg/mL BSA. To prepare pHrodo-labeled TGT beads, TGT beads were incubated with pHrodo-SE (200 ng/mL; P36600, Invitrogen) at 25°C for 30 min, washed, and resuspended in DMEM containing 5 mg/mL BSA. To prepare TGT plates, PEGylated coverslips (PEG-biotin:PEG=1:40, 26 mm × 40 mm) were immobilized on a 60 mm Petri dish using epoxy. The PEGylated coverslips were dotted with 1.6 μL neutravidin (200 μg/mL) and incubated at 25°C for 20 min. Then, the coverslips were tilted by hand, washed with cold phosphate-buffered saline (PBS), dotted with 1 μL of 5 μM TGT, and incubated at 4°C for 30 min. Finally, TGT plates were washed twice with PBS without drying.

**Phagocytosis assay**

The phagocytosis assay was performed as previously described (*17*). Briefly, phagocytes were plated on a 24-well culture plate, incubated with TGT beads or apoptotic cells at 37°C in serum-free medium, extensively washed with ice-cold PBS, trypsinized, and analyzed by flow cytometry or confocal microscopy. For the *in vivo* phagocytosis assay, TAMRA-labeled apoptotic Jurkat cells were treated with or without cytochalasin D (2 μM), washed, and resuspended in Dulbecco's PBS (DPBS). A total of 10$^7$ apoptotic Jurkat cells in 300 μL DPBS were intraperitoneally injected into 8-week-old mice. At 30 min after the injection, the mice were sacrificed and peritoneal exudates were collected. Exudate cells were stained with an anti-mouse F4/80 antibody and analyzed by flow cytometry. For the real-time phagocytosis assay, J774A.1 cells stably expressing Raichu-Rac1, PH-Akt-GFP, or membrane-targeted GFP plated on a 35 mm confocal dish were incubated with TGT beads. F-actin was stained with 5 μM SiR-actin (CY-SC001, Cytoskeleton) for 30 min at 37°C before incubation with TGT beads. The cells were observed by time-lapse confocal microscopy.

**Frustrated phagocytosis assay**

Phagocytes ($10^6$/mL) suspended in serum-free DMEM were plated on TGT plates, incubated in a $CO_2$ incubator at 37°C for 10 or 60 min, washed with PBS, and observed by microscopy. For the real-time frustrated phagocytosis assay, the pre-attached glass of a 35 mm confocal dish was removed and a PEGylated coverslip (PEG-biotin:PEG=1:40, 22 mm × 22 mm) was immobilized using epoxy. The subsequent process was the same as that used to prepare TGT plates. J774A.1 cells stably expressing membrane-targeted GFP (2 × $10^5$/mL) were plated on PS-TGT plates and observed by real-time microscopy. To observe rupture of the TGT, phagocytes were plated on plates coated with a 1:1 mixture of Cy5-labeled PS-$TGT^{56pN}$ or PS-$TGT^{12pN}$ and Cy3-labeled PS-$TGT^{56pN}$. Fluorescence signals were observed by confocal microscopy after incubation in a $CO_2$ incubator at 37°C for 60 min.

**Platelet preparation**

Mice were sacrificed at 11–15 weeks of age. Then, nine volumes of blood were collected in one volume of ACD solution at 37°C by cardiac puncture. Platelet-rich plasma was collected by centrifugation at 200 × g for 20 min, mixed with an equal volume of modified Tyrode's buffer (12 mM $NaHCO_3$, 68.44 mM NaCl, 5.55 mM D-glucose, 2.24 mM $MgCl_2$, 2.68 mM KCl, 2.38 mM $CaCl_2$, 0.41 mM $NaH_2PO_4$, and 10 mM HEPES, pH 7.4) containing 0.3 μM PGE1 and 15% (vol/vol) ACD, and centrifuged at 1,000 × g for 5 min. Pelleted platelets were washed with modified Tyrode's buffer supplemented 0.15 μM PGE1 and 1 mM EDTA and resuspended in an adequate volume of modified Tyrode's buffer.

**Generation of PS-exposing live cells**

Jurkat cells were labeled with 25 μM TAMRA-SE (C1171, Invitrogen) for 15 min, washed, exposed to 100 mJ/$cm^2$ UV irradiation, and incubated in a $CO_2$ incubator at 37°C for 3 h. To modulate membrane tension, apoptotic Jurkat cells were treated with cytochalasin D (2 μM) at

37°C for 1 h and then extensively washed with RPMI. PS exposure was confirmed by propidium iodide (PI)-Annexin V staining. To induce PS exposure on live Jurkat cells, TAMRA-SE-labeled Jurkat cells were incubated in serum-free RPMI containing 1 mM MβCD at 37°C for 30 min. Thereafter, the cells were washed with serum-free RPMI and incubated in calcium-free DPBS containing 10 μM A23187 at 25°C for 15 min to induce PS exposure. Platelets were labeled with 10 μM CellTracker Green CMFDA (C7025, Invitrogen) for 20 min at 37°C in modified Tyrode's buffer supplemented with 0.15 μM PGE1 and washed once with modified Tyrode's buffer containing 0.05% BSA. Platelets were activated by treatment with 0.5 U/mL thrombin and 10 μg/mL collagen at 25°C for 15 min and further incubated with or without 10 mM MβCD at 25°C for 30 min.

**Immunofluorescence staining**

J774A.1 cells were plated on a 35-mm confocal dish, incubated with TGT beads in a $CO_2$ incubator at 37°C for 15 min, washed five times with PBS at 4°C, and fixed with 4% paraformaldehyde diluted in PBS at 25°C for 15 min. Thereafter, the cells were permeabilized by treatment with 0.1% Triton X-100 for 5 min and blocked by treatment with 10% BSA for 30 min. The cells were then incubated with an anti-MLC (phospho S20) antibody (1:100), an anti-Tim-4 antibody (1:50), an anti-PI3K antibody (1:100), or Alexa Fluor 594-conjugated phalloidin (1:20) diluted in PBS containing 3% BSA overnight at 4°C, stained with an Alexa Fluor 488-conjugated goat anti-mouse antibody (1:2000), an Alexa Fluor 647-conjugated donkey anti-rabbit antibody (1:2000), or an Alexa Fluor 488-conjugated goat anti-rabbit antibody (1:2000) for 1 h, and finally incubated with Hoechst 33342 (1:10000) diluted in PBS for 10 min. Images were acquired using an Olympus FV1000 SPD system (Olympus, Tokyo, Japan).

**Immunoblotting and immunoprecipitation**

293T, Jurkat, or J774A.1 cells were lysed using lysis buffer (50 mM Tris (pH 7.6), 150 mM NaCl,

10 mM NaPP, 10 mM NaF, 1 mM $Na_3VO_4$, 1% Triton X-100, 10 µg/mL pepstatin, 10 µg/mL leupeptin, 10 µg/mL AEBSF, and 10 µg/mL aprotinin). Proteins were separated by SDS-PAGE, transferred to a nitrocellulose membrane, and detected using appropriate antibodies. For immunoprecipitation, lysates were incubated with anti-HA antibody-conjugated agarose beads or with an anti-PI3K antibody and protein A/G agarose beads (sc-2003, Santa Cruz) for 2 h. Proteins bound to the beads were separated by SDS-PAGE, transferred to a nitrocellulose membrane, and detected by immunoblotting.

**Analysis of Rac1 activity during phagocytosis**

*Unlabeled (differential interference contrast (DIC)) bead detection*: A reference z-stack DIC image of a single bead was prepared prior to the experiment. The bead image obtained during phagocytosis of TGT beads by J774A.1 cells stably expressing Raichu-Rac1 was first analyzed using the circular Hough transform to determine the rough position of the beads. Starting from the initial estimated position, the reference image set was least-square fitted with the bead image by interpolating the reference image along four parameters (bead position x, y, z and radius r). If two detected points were less separated than their radii, one point with lower radial symmetry (ratio of the standard deviation and average of the angular intensity profile) was rejected.

*Cell segmentation*: YFP images of J774A.1 cells stably expressing Raichu-Rac1 were filtered with 1 pixel neighbor(8-connectivity) 2d median filtering to reduce noise. The intensity dynamic range was adjusted to saturate 5% of the highest and lowest intensities. A Tophat filter and dilation with a 7 pixel disk were applied to remove detailed features and ensure the connectivity of segments from the same cell. To determine the threshold intensity of each cell, multiple reference regions (21×21 pixels) of each cell were manually selected, and the processed image was thresholded by the minimum intensity in these reference regions. Connected components of this thresholded image were obtained, and only connected components containing the reference region were selected and assigned for each cell.

*FRET ratio calculation*: The respective background intensities calculated by the mean intensity value of pixels inside the background mask were subtracted from CFP and YFP images. The 90% quantile thresholded image of total intensity (CFP + YFP intensities) was used as a background mask. Each image was filtered with 1 pixel neighbor(8-connectivity) 2d median filtering to remove noise. The FRET ratio was calculated as the ratio between the processed YFP and CFP images. It was presented in a heatmap by mapping the FRET ratio using color and the total intensity using opacity. The FRET ratio and total intensity were collected for every cell, and the representative FRET value for each cell was calculated as the total intensity weighted mean of the FRET ratio. Every analysis was performed using MATLAB custom code.

**Analysis of PI3K activity during phagocytosis**

*Labeled bead detection*: The image of fluorescence surface-labeled beads, which had a donut-like shape, was analyzed by a custom method similar to the circular Hough transform. Five images of convolution kernels were built, in which the radial intensity profile was gaussian with offset (40–80 pixels), to model the donut-like images of beads of various sizes. Each kernel was used to convolute a bead image, and the pixel-wise maximum image M among convoluted images was calculated. M was thresholded by over both 70% of the maximum intensity and gaussian filtered (sigma=60 pixels) image of M/0.4. The thresholded image T was denoised by 1 pixel neighbor(8-connectivity) 2 d median filtering and dilation (2 pixel disk). The connected component of T was identified and its center of mass (CM) was calculated. CMs with radial symmetry less than 0.6 were rejected. GFP intensities surrounding the beads were collected to determine the PH-AKT concentration.

*Cell segmentation*: GFP images of J774A.1 cells stably expressing PH-Akt-GFP were first analyzed to determine the background level. A median filter (5×5 pixel window), 1% extrema saturation, dilation (2 pixel disk), gaussian blur (1 pixel), and thresholding (intensity lower than

Otsu’s threshold × 0.5) were applied to set the background region. Then, the GFP image was thresholded using this background level × 1.05, and the cytosol region and the membrane region were obtained and rejected using a 10 pixel disk kernel. The average intensity of this region, excluding the bead-contacting region, was collected and used as the cytosol intensity. Every analysis was performed using MATLAB custom code.

**Proximity ligation assay**

Proximal ligation assays were performed according to the manufacturer's protocol (DUO92002; DUO92004; DUO92008, Sigma Aldrich, St. Louis, MO, USA). Peritoneal macrophages were plated on 18 mm-diameter cover glasses, incubated with Cy3-labeled PS-TGT beads for 10 min. Then, peritoneal macrophages were fixed with 4% paraformaldehyde, permeabilized by treatment with 0.1% Triton X-100, blocked with blocking solution, and incubated with anti-TIMD4 C-term antibody (RB40431, Raybiotech) and anti-PI3K (ab86714, Abcam) antibody at 4°C overnight. After that, peritoneal macrophages were incubated with anti-rabbit PLUS probe and anti-mouse MINUS probe, then incubated with amplification buffer. Images were observed using Olympus FV3000RS confocal microscopy (Olympus, Tokyo, Japan).

**Synthesis of Hairpin PS-TGT (PEG loop)**

The upper strand was conjugated with PS as previously described for PS-TGT, except the Cy3-NHS labeling, leaving amine groups unmodified. The extended lower strand had thiol and amine added for fluorophore labeling and loop conjugation, respectively, and was purchased from Integrated DNA Technologies (12 pN: 5’-/5Biosg/GTG TCG TGC CTC CGT GCT GTG TT/iAmMC6T/ TT/3ThioMC3-D/-3’, 56 pN: 5’-GTG TCG TGC CTC CGT GCT GTG /iBiodT/T/iAmMC6T/ TT/3ThioMC3-D/-3’). These lower strands were labeled with Cy3-NHS as previously described, precipitated with ethanol, and dissolved in PBS with 3.3 mM TCEP to a final concentration of 167 μM. This solution was degassed under vacuum for 20 min to facilitate thiol deprotection. Subsequently, a 1:1.4 molar ratio of the PS-conjugated upper strand and a 1:2 molar ratio of Maleimide-PEG-SVA (molecular weights of 2, 5, 10, and 20 kDa, Laysan Bio) were added, and incubated overnight at 25°C. The correct-sized products were then extracted using PAGE gel extraction.

**Synthesis of Hairpin PS-TGT (DNA loop)**

8 nt loop fragment was concatenated to the 3' end of upper strand (5'-/5ThioMC6-D/CAC AGC ACG GAG GCA CGA CAC G/iAmMC6T/CT ACC C-3') and 5' end of lower strand with phosphate group (12 pN: 5'-/5Phos/ATC TAA GTG TGT CGT GCC TCC GTG CTG TG/3Bio/-3', 56 pN: 5'-/5Phos/ATC TAA G/iBiodT/GT GTC GTG CCT CCG TGC TGT G-3'), in order to have sticky end for ligation with loop DNA. Conjugation of PS and Cy3 to the upper strand was performed as previously described method for PS-TGT. 1:2 molar ratio of loop strand (50 nt DNA loop: 5'-/5Phos/AAT AAA GTT TAT CTT TTT TTT TGT TTG AAT ACC C-3', 100 nt DNA loop: 5'-/5Phos/AAT AAA GTT TAT C ($T_{59}$) GTT TGA ATA CCC-3') and splint strand (splint for upper-loop: 5'-/5Phos/GAT AAA CTT TAT TGG GTA GA-3', splint for lower-loop: 5'-/5Phos/CTT AGA TGG GTA TTC AAA CA-3') was added. For 16 nt DNA loop, only splint strand was added (5'-/5Phos/CTT AGA TGG GTA GA-3') to directly ligate loop fragments included in the upper and lower strands. Ligation was then performed using 4.2 U/pmol of T4 ligase (M0202S, New England Biolabs). To remove the splint strand, a 20-fold excess of the complement strand was added as a trap, followed by heating up to 95°C for 3 min and slow cooling.

**Statistical analysis**

The sample numbers (n) are indicated in the figure legends. All data except box-and-whisker plots are presented as mean ± s.e.m. In box-and-whisker plots, the horizontal lines of the box represent the median along with the first and third quartiles, and the whiskers indicate the data range from minimum to maximum. The two-tailed unpaired Student's t test, a one-way ANOVA, or a two-way ANOVA were used to analyze statistical differences. Analysis was performed using GraphPad Prism 6 software.

### Supplementary Text

This Supplementary Text includes uncropped blots of western blot panels, the gating strategies used for flow cytometry analysis, movie legends, and synthesis of 18:1-12:0(maleimide) phosphatidylcholine.

### Uncropped blots of western blot panels

#### Fig. 3D

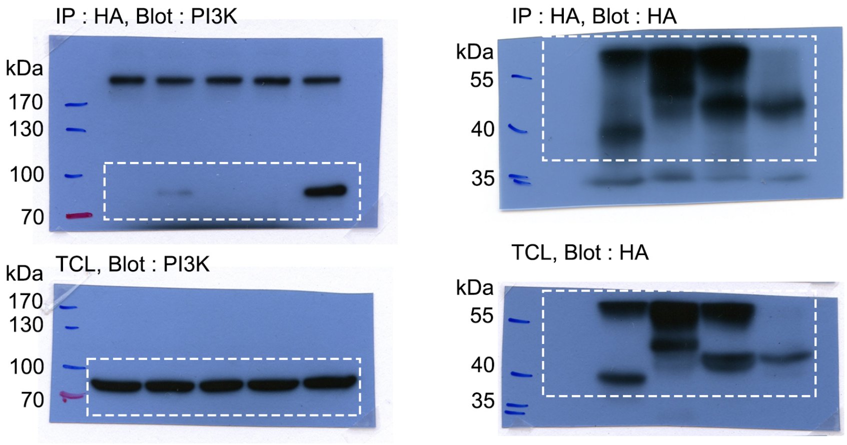


#### Fig. S4A

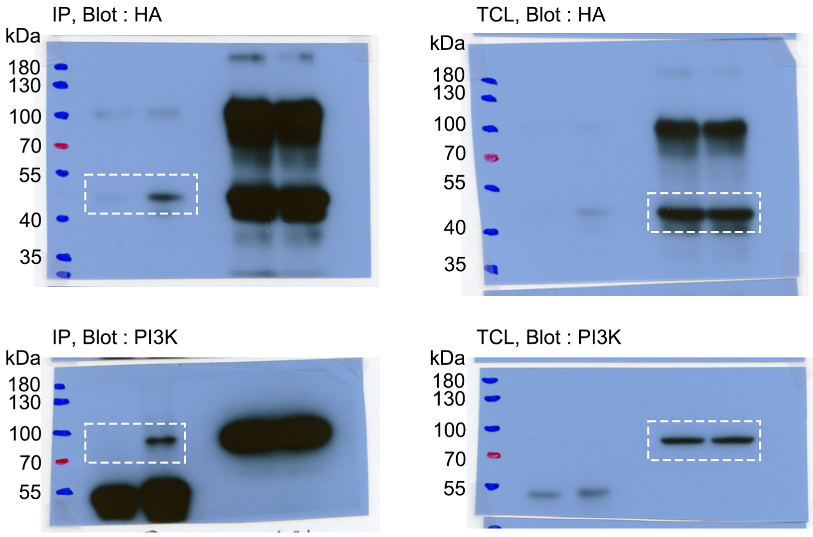

#### Fig. S4B

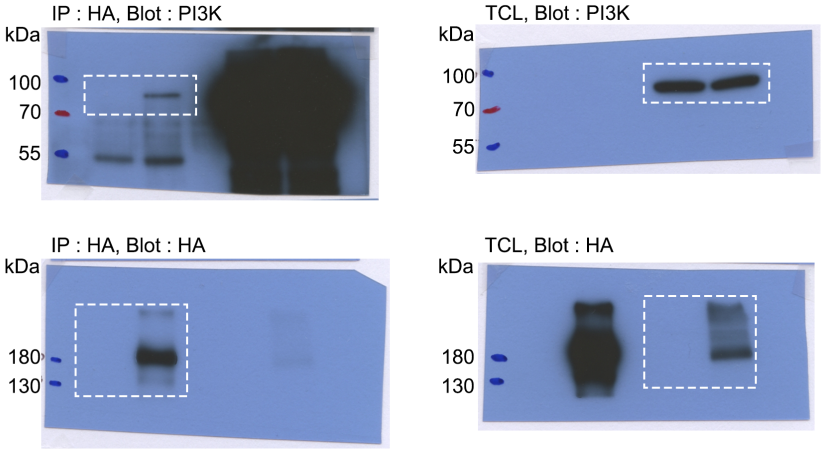


#### Fig. S5E

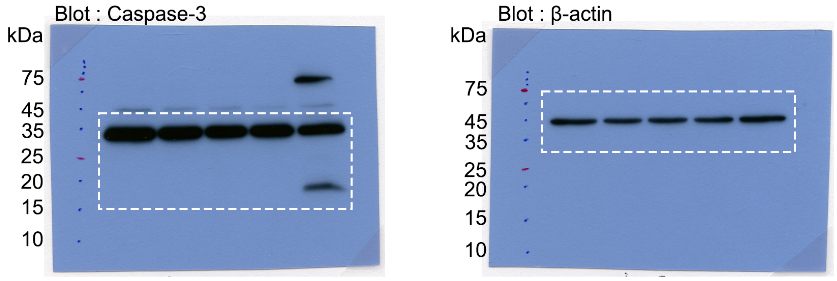


**Uncropped blots for western blot panels presented**

Uncropped blots for western blot panels presented are outlined. Figure panels are indicated above the corresponding western blot set. White dashed lines indicate where membranes were cut.

### Gating strategies used for flow cytometry analysis

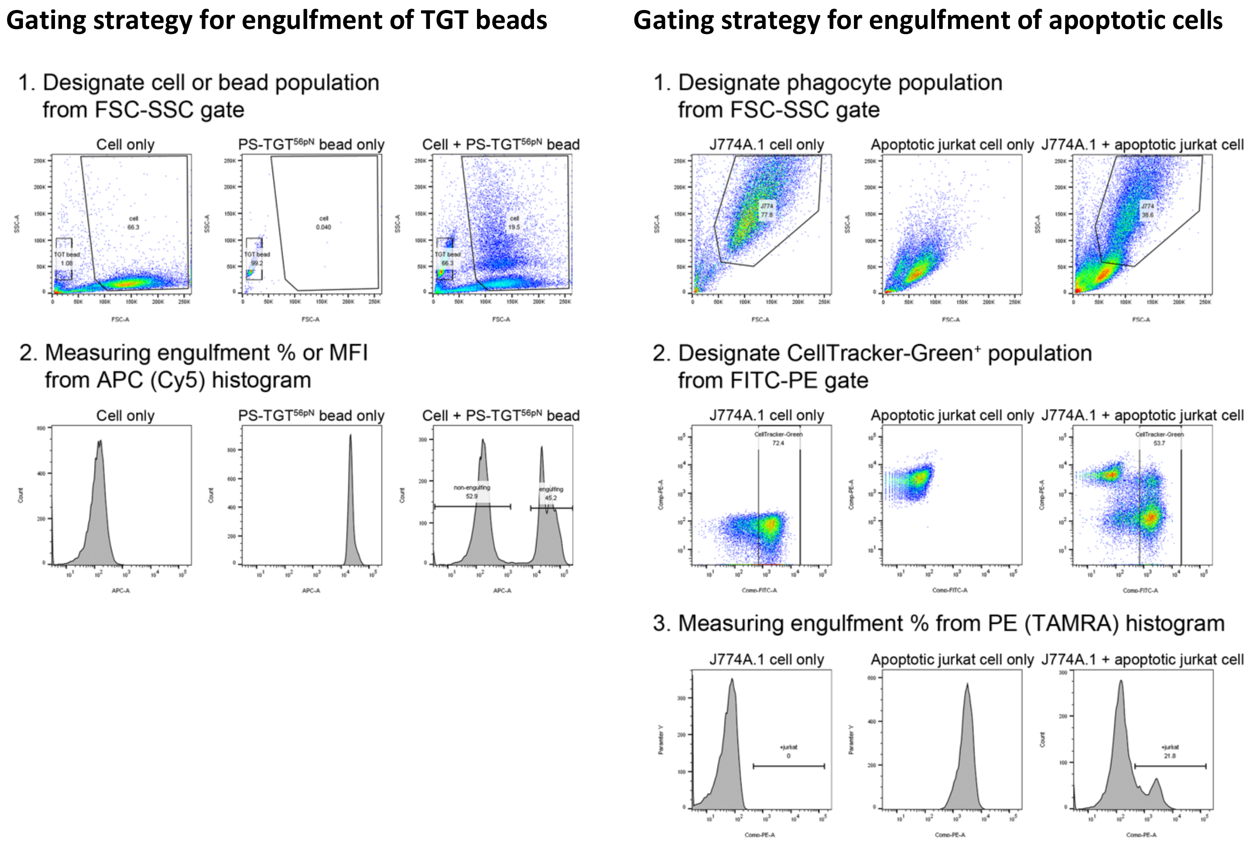


##### Gating strategies for flow cytometry analysis

Gating strategies for flow cytometry analysis of engulfment of TGT beads (left) or apoptotic cells (right) presented in the manuscript are outlined.

#### Movie legends

**Movie S1. Engulfment of PS-TGT$^{12pN}$ and PS-TGT$^{56pN}$ beads by J774A.1 cells**
A 30 min time-lapse video starting at addition of the indicated Cy3-labeled PS-TGT beads. J774A.1 cells were incubated with Cy3-labeled PS-TGT beads and observed by confocal microscopy. Images were obtained every 20 sec for 30 min. The cells in rectangles are shown in Fig. 1B. Scale bar, 20 μm.

**Movie S2. Phagolysosomal acidification during engulfment of the indicated PS-TGT beads** A 30 min time-lapse video starting at addition of the indicated Cy5- and pHrodo-labeled PS-TGT beads. J774A.1 cells stably expressing membrane-targeted GFP were incubated with Cy5- and pHrodo-labeled PS-TGT beads and observed by confocal microscopy. Images were obtained every 20 sec for 30 min. Scale bar, 10 μm.

**Movie S3. Analysis of pHrodo intensities during engulfment of PS-TGT beads**
A representative example video for pHrodo intensity analysis during engulfment of PS-TGT beads. MATLAB was used to analyze the pHrodo intensity of PS-TGT$^{56pN}$ beads after they had been completely ingested. A 30 min time-lapse video of GFP and pHrodo (upper left) and GFP and Cy5 (upper right). Red-circled and cyan-circled beads indicate unengulfed and completely engulfed beads, respectively (upper right). The pHrodo intensity of each bead was tracked in a different color. The thick black line indicates the average of pHrodo intensities (lower).

**Movie S4. F-actin duration during engulfment of the indicated PS-TGT beads**
A time-lapse video starting at addition of the indicated Cy3-labeled PS-TGT beads. J774A.1 cells were stained with CellTracker and SiR-actin, incubated with the indicated PS-TGT beads, and observed by confocal microscopy. Images were obtained every 20 sec. Scale bar, 10 μm.

**Movie S5. Phagocytic cup closure during engulfment of PS-TGT beads**
A 30 min time-lapse video starting at addition of the indicated PS-TGT beads. J774A.1 cells stably expressing membrane-targeted GFP were incubated with the indicated PS-TGT beads and observed by confocal microscopy. Images were obtained every 20 sec for 30 min. Scale bar, 10 μm.

**Movie S6. Rac1 activity during engulfment of PS-TGT beads**

A 30 min time-lapse video starting at addition of the indicated PS-TGT beads. J774A.1 cells stably expressing Raichu-Rac1 were incubated with the indicated PS-TGT beads and observed by confocal microscopy. Images were obtained every 20 sec for 30 min. Scale bar, 10 μm.

**Movie S7. Rac1 activity analysis during engulfment of PS-TGT beads**

A representative example video for Rac1 activity analysis during engulfment of PS-TGT beads. MATLAB was used to analyze Rac1 activity in the phagocytic cup during engulfment of PS-$TGT^{56pN}$ beads. A 30 min time-lapse merged video of DIC and Rac1-FRET (upper left). Red-circled and cyan-circled beads indicate unengulfed and completely engulfed beads, respectively. The initial point of contact between the phagocyte and the bead was aligned to 0° of the phagocytic cup angle (lower left). Two-dimensional (upper right) and three-dimensional (lower right) time-lapse videos for Rac1 activity analysis in the phagocytic cup.

**Movie S8. Expansion and contraction of phagocytes on PS-TGT plates**

A time-lapse video starting at plating of J774A.1 cells. J774A.1 cells stably expressing membrane-targeted GFP were incubated on the indicated PS-TGT plates and observed by confocal microscopy. Images were obtained every 20 sec. Scale bar, 10 μm.

**Movie S9. Effect of blebbistatin on expansion and contraction of phagocytes on PS-TGT plates**

A 30 min time-lapse video starting at plating of J774A.1 cells in the presence of blebbistatin. J774A.1 cells stably expressing membrane-targeted GFP were incubated on the indicated PS-TGT plates in the presence of 10 μM blebbistatin and observed by confocal microscopy. Images were obtained every 20 sec for 30 min. Scale bar, 10 μm.

**Movie S10. Area analysis of J774A.1 cells on PS-TGT plates**

A representative example video for analysis of the normalized area of J774A.1 cells on PS-TGT plates. MATLAB was used to analyze the area changes of J774A.1 cells incubated on a PS-$TGT^{56pN}$ plate. J774A.1 cells stably expressing membrane-targeted GFP on a PS-$TGT^{56pN}$ plate were observed by confocal microscopy (left). GFP-positive regions were considered the areas where cells spread, and each cell is represented in a different color (middle). The normalized area of each cell is expressed in a different color, and the average value is indicated by a thick black line (right).

**Movie S11. Phosphorylation of MLC in J774A.1 cells on PS-TGT plates**

Three-dimensional images of pMLC in J774A.1 cells on PS-TGT plates. J774A.1 cells were incubated on PS-TGT plates for 60 min, washed, fixed, stained with an anti-pMLC antibody and phalloidin, and observed by confocal microscopy. Scale bar, 20 μm.

**Movie S12. PI3K activity during engulfment of PS-TGT beads**

A time-lapse video starting at addition of the indicated PS-TGT beads. J774A.1 cells stably expressing PH-Akt-GFP were incubated with the indicated Cy3-labeled PS-TGT beads and observed by confocal microscopy. Images were obtained every 20 sec. Scale bar, 10 μm.

**Movie S13. The effect of LY294002 on PI3K activity during engulfment of PS-TGT beads**

A 30 min time-lapse video starting at addition of the indicated PS-TGT beads in the presence of LY294002. J774A.1 cells stably expressing PH-Akt-GFP were incubated with the indicated Cy3-labeled PS-TGT beads in the presence of 50 μM LY294002 and observed by confocal microscopy. Images were obtained every 20 sec for 30 min. Scale bar, 10 μm.

**Movie S14. PI3K activity analysis during engulfment of PS-TGT beads**

A representative example video for PI3K activity analysis during engulfment of PS-TGT beads. MATLAB was used to analyze PI3K activity during engulfment of PS-TGT$^{56pN}$ beads. A 30 min time-lapse video for PH-Akt-GFP and Cy3 (lower left). The membrane and cytosol of a phagocyte engulfing beads are marked in magenta and green, respectively, and each bead being engulfed is represented by a different color (upper left). The GFP intensity in the phagocytic cup relative to that in the cytosol is dotted in a different color, and the average relative GFP intensity is indicated by a thick black line (upper right). Time-lapse changes of the GFP intensity in the phagocytic cup relative to that in the cytosol are expressed in a parula colormap (lower right).

**Movie S15. The effect of hypertonic medium on PI3K activity during engulfment of PS-TGT$^{56pN}$ beads**

A time-lapse video starting at addition of the indicated PS-TGT$^{56pN}$ beads. J774A.1 cells stably expressing PH-Akt-GFP were incubated with Cy3-labeled PS-TGT$^{56pN}$ beads in control or hypertonic medium and observed by confocal microscopy. Images were obtained every 20 sec. Scale bar, 10 μm.

### Synthesis of 18:1-12:0(maleimide) phosphatidylcholine

##### 1. General Information of 18:1-12:0(maleimide) PC synthesis

Reactions were performed in oven-dried (140°C) or flame-dried glassware under an atmosphere of dry argon unless otherwise noted. Dichloromethane ($CH_2Cl_2$) and *N*,*N*-dimethylformamide (DMF) were dried by percolation through a column packed with neutral alumina and a column packed with Q5 reactant, a supported copper catalyst for scavenging oxygen, under a positive pressure of argon. Solvents used for workup and chromatography were acetone (Duksan, Extra Pure grade), ethanol (Duksan, Extra Pure grade), dichloromethane (Duksan, Extra Pure grade), and methanol (Fisher, HPLC grade). Silica gel 60 ($SiO_2$, Merck Millipore, 40–63 μm) and alumina 60 ($Al_2O_3$, Acros, neutral, Brockmann I, 50–200 μm) were used for column chromatography and filtration. Merck Millipore TLC silica gel 60 $F_{254}$ was used for thin layer chromatography (TLC).

1,8-Diazabicyclo[5.4.0]undec-7-ene (DBU, Alfa, $CaH_2$) and *N*-methylimidazole (NMI, Alfa, Na) were distilled over indicated drying agents under reduced pressure before use. Diisopropylethylamine (DIPEA) was distilled over $CaH_2$ under argon before use. The following reagents were used as received: lecithin (Soybean, Alfa, 90%), 2-methyl-6-nitrobenzoic anhydride (MNBA, Aldrich, 97%), *N*,*N'*-dicyclohexylcarbodiimide (DCC, TCI, 98%), 4-(dimethylamino)pyridine (DMAP, Alfa, 99%), 1-[bis(dimethylamino)methylene]-1*H*-1,2,3-triazolo[4,5-*b*]pyridinium 3-oxide hexafluorophosphate (HATU, Chem-Impex, 99.9%), (benzotriazol-1-yloxy)tripyrrolidinophosphonium hexafluorophosphate (pyBOP, Novabiochem, 99%), and 1-hydroxybenzotriazole hydrate (HOBt, Aldrich, 97%).

$^{1}H$ and $^{13}C$ NMR spectra were recorded on a Jeol ECS400 spectrometer (400 MHz, $^{1}H$; 100 MHz, $^{13}C$; 162 MHz, $^{31}P$). $^{1}H$ and $^{13}C$ NMR spectra were referenced to residual chloroform (7.26 ppm, $^{1}H$; 77.23 ppm, $^{13}C$). $^{31}P$ NMR spectra were recorded using triphenyl phosphate as an internal standard (δ –17.70 ppm in $CDCl_3$). Chemical shifts are reported in ppm, and multiplicities are indicated by s (singlet), d (doublet), t (triplet), q (quartet), br (broad), and m (multiplet). Coupling constants, *J*, are reported in Hertz. High resolution mass spectrometry (HRMS) was performed on an Agilent 6520 Q-TOF mass spectrometer and a Thermo Scientific LTQ Orbitrap XL mass spectrometer at Environmental OMICS Laboratory, GIST. The mass spectral data are reported in the form of (m/z). UV (254 nm), a potassium permanganate ($KMnO_4$) staining solution, and a ceric ammonium molybdate (CAM) staining solution were used for visualization of TLC.

*sn*-Glycero-3-phosphocholine (**1**),[1] 1-oleoyl-*sn*-glycero-3-phosphocholine (**2**),[2] 12-[*N*-(9-fluorenylmethoxycarbonyl)amino]dodecanoic acid (**3**),[3] and 4-(*N*-maleimidomethyl)cyclohexanecarboxylic acid (**4**)[4] were prepared according to the cited literature procedures.

#### 2. Experimental Procedures

##### 2.1. Isolation of Phosphatidylcholines from Soybean Lecithin

Phosphatidylcholines were prepared according to the patent process with modification.[5] Soybean lecithin was suspended in hexanes and heated until the mixture turned dark brown gummy solid. The thermally altered gum was suspended in acetone and stirred vigorously overnight. The resulting mixture was filtered to give brown solid, which was suspended in ethanol and stirred vigorously overnight. The ethanol-soluble fraction was isolated by celite filtration and stirred again with alumina. The resulting mixture was filtered through celite and concentrated under reduced pressure to give phosphatidylcholines as a yellow sticky oil, which was used for the preparation of *sn*-glycero-3-phosphocholine (**1**) without further purification.

##### 2.2. Acylation of 1-Oleoyl-*sn*-glycero-3-phosphocholine (2)[6]

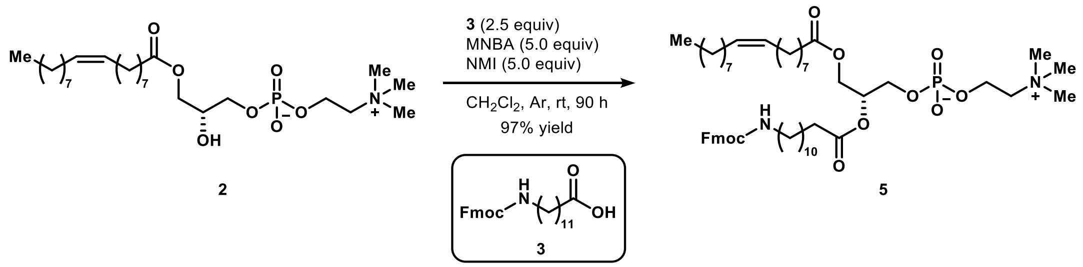


To a mixture of **3** (95 mg, 0.18 mmol) and NMI (73 μL, 0.91 mmol) in $CH_2Cl_2$ (1 mL) was added dropwise a solution of MNBA (317 mg, 0.920 mmol) in $CH_2Cl_2$ (4.6 mL) at room temperature under argon. After 0.5 h, a solution of **2** in $CH_2Cl_2$ (4 mL) was added to the reaction mixture. After being stirred for 90 h, the reaction mixture was concentrated under reduced pressure. The crude mixture was purified by column chromatography ($SiO_2$, $\phi$ = 3.0 cm, $l$ = 8.0 cm, $CH_2Cl_2$:MeOH= 3:1 to $CH_2Cl_2$:MeOH:$H_2O$ = 18:6:1, $R_f$ = 0.68 in $CH_2Cl_2$:MeOH:$H_2O$ = 18:6:1, CAM) to afford **5** as a white sticky oil (167 mg, 97% yield).

Data for **5**: JSJ-11-025

$^1$H NMR (400 MHz, $CDCl_3$) δ 7.76 (d, *J* = 7.6, 2H), 7.59 (d, *J* = 7.6, 2H), 7.39 (t, *J* = 7.3, 2H), 7.30 (td, *J* = 7.5, 1.0, 2H), 5.38–5.29 (m, 2H), 5.25–5.18 (m, 1H), 4.86–4.78 (m, 1H), 4.46–4.34 (m, 5H), 4.24–4.18 (m, 1H), 4.13 (dd, *J* = 12.1, 7.2, 1H), 4.05–3.98 (m, 2H), 3.95–3.89 (m, 2H), 3.39 (s, 9H), 3.22–3.14 (m, 2H), 2.30 (t, *J* = 7.5, 2H), 2.28 (t, *J* = 7.9, 2H), 2.05–1.96 (m, 4H), 1.64–1.53 (m, 4H), 1.53–1.44 (m, 2H), 1.37–1.20 (m, 34H), 0.87 (t, *J* = 6.9, 3H).

$^{13}$C NMR (100 MHz, $CDCl_3$) δ 173.7, 173.4, 156.6, 144.2 (2C), 141.4 (2C), 130.2, 129.9, 127.8 (2C), 127.2 (2C), 125.2 (2C), 120.1 (2C), 70.7 (d, *J* = 6.7), 66.7, 66.6 (d, *J* = 3.8), 63.6, 63.2, 59.5 (d, *J* = 3.8), 54.6 (3C), 47.5, 41.3, 34.5, 34.3, 32.1, 30.2–29.3 (15C), 27.41, 27.39, 27.0, 25.12, 25.07, 22.9, 14.3.

$^{31}$P NMR (162 MHz, $CDCl_3$) δ –1.43.

HRMS (ESI) m/z calcd for $C_{53}H_{86}N_2O_{10}P$ $[M + H]^+$ 941.6015, found 941.6008.

#### 2.3. In Situ Fmoc-Deprotection/Amide Formation of 5

##### Table S1. Survey of coupling reagents

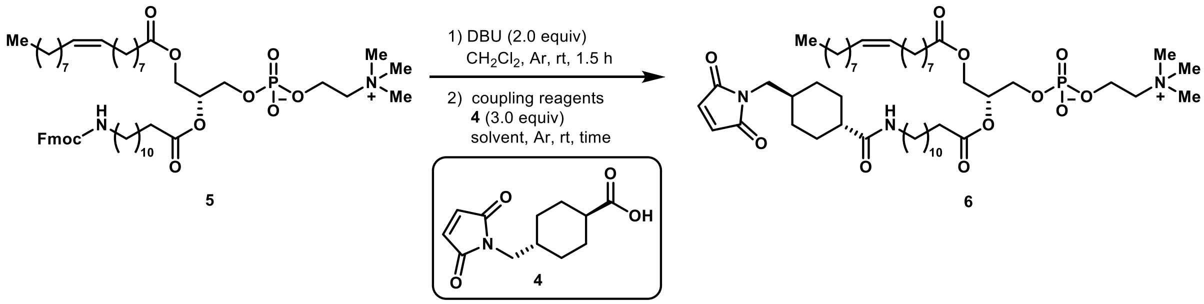


| entry | coupling reagents | solvent | time (h) | yield (%) |
|---|---|---|---|---|
| 1 | MNBA (4.5 equiv), NMI (4.5 equiv) | $CH_2Cl_2$ | 3 | 72 |
| 2 | HATU (3.0 equiv), DIPEA (6.0 equiv) | DMF | 17.5 | 100 |
| 3 | pyBOP (3.0 equiv), HOBt (3.0 equiv) DIPEA (6.0 equiv) | DMF | 17.5 | 84 |
| 4 | DCC (3.0 equiv), DMAP (0.5 equiv) | $CH_2Cl_2$ | 18 | 0 |

Several coupling reagents were examined for amide formation (Table S1). The desired product was obtained in good yields by employing MNBA, HATU, and pyBOP/HOBt (entries 1–3). On the other hand, a complex mixture was produced from the reaction with DCC/DMAP (entry 4).

**General procedure for Table S1, entries 1–3**

A solution of **5** (19 mg, 0.020 mmol) and DBU (6 μL, 0.04 mmol) in $CH_2Cl_2$ (1 mL) was stirred at room temperature under argon for 1.5 h for deprotection of the Fmoc group. The resulting mixture was added to a solution of **4** (14 mg, 0.060 mmol) and coupling reagents in $CH_2Cl_2$ (0.9 mL) or DMF (1.2 mL) at room temperature under argon. After being stirred for the time shown in Table S1, the reaction mixture was concentrated under reduced pressure. The crude mixture was purified by column chromatography ($SiO_2$, $\phi$ = 1.5 cm, $l$ = 8.0–10.0 cm, $CH_2Cl_2$:MeOH= 3:1 to $CH_2Cl_2$:MeOH:$H_2O$ = 18:6:1, $R_f$ = 0.63 in $CH_2Cl_2$:MeOH:$H_2O$ = 18:6:1, CAM) to afford **6** as a white sticky oil which was contaminated by a small amount of unidentified impurities.

The products from the three reactions were combined and purified again by column chromatography ($SiO_2$, $\phi$ = 2.5 cm, *l* = 11.0 cm, $CH_2Cl_2$:MeOH= 1:1 to $CH_2Cl_2$:MeOH:$H_2O$ = 90:30:3, $R_f$ = 0.53 in $CH_2Cl_2$:MeOH:$H_2O$ = 90:30:3, CAM) to afford **6** as a white sticky oil with improved purity (29 mg, 52% yield).

<u>Data for **6**</u>: JSJ-11-028, 030, 031

$^1$H NMR (400 MHz, $CDCl_3$) δ 6.70 (s, 2H), 5.59–5.53 (m, 1H), 5.39–5.30 (m, 2H), 5.26–5.18 (m, 1H), 4.41 (dd, *J* = 12.4, 2.6, 1H), 4.40–4.32 (m, 2H), 4.13 (dd, *J* = 12.1, 6.9, 1H), 4.02–3.96 (m, 2H), 3.84–3.79 (m, 2H), 3.39 (s, 9H), 3.36 (d, *J* = 7.0, 2H), 3.24–3.17 (m, 2H), 2.29 (t, *J* = 7.5, 2H), 2.28 (t, *J* = 7.5, 2H), 2.08–1.95 (m, 5H), 1.92–1.84 (m, 2H), 1.77–1.65 (m, 4H), 1.63–1.52 (m, 4H), 1.49–1.39 (m, 5H), 1.36–1.22 (m, 32H), 1.05–0.96 (m, 2H), 0.88 (t, *J* = 6.9, 3H).

$^{13}$C NMR (100 MHz, $CDCl_3$) δ 175.7, 173.8, 173.5, 171.3 (2C), 134.2 (2C), 130.2, 129.9, 70.8 (d, *J* = 7.3), 66.8 (d, *J* = 5.0), 63.6 (d, *J* = 5.0), 63.2, 59.4 (d, *J* = 3.2), 54.8 (3C), 45.5, 43.9, 39.6, 36.6, 34.5, 34.3, 32.1, 30.1–29.1 (19C), 27.44, 27.40, 27.1, 25.12, 25.09, 22.9, 14.3.

$^{31}$P NMR (162 MHz, $CDCl_3$) δ –0.65.

HRMS (ESI) m/z calcd for $C_{50}H_{89}N_3O_{11}P$ $[M + H]^+$ 938.6229, found 938.6238.

#### 3. References

1. Park, J. M.; De Castro, K. A.; Ahn, H.; Rhee, H. *Bull. Korean Chem. Soc.* **2010**, *31*, 2689–2691.

2. Pierrat, P.; Kereselidze, D.; Lux, M.; Lebeau, L.; Pons, F. *Int. J. Pharm.* **2016**, *511*, 205–218.

3. Chen, W.-H.; Chen, J.-X.; Cheng, H.; Chen, C.-S.; Yang, J.; Xu, X.-D.; Wang, Y.; Zhuo, R.-X.; Zhang, X.-Z. *Chem. Commun.* **2013**, *49*, 6403–6405.

4. Christie, R. J.; Anderson, D. J.; Grainger, D. W. *Bioconjugate Chem.* **2010**, *21*, 1779–1787.

5. Weete, J. D.; Griffith, G. L. Process for Obtaining Highly Purified Phosphatidylcholine. U.S. Patent 5,453,523, September 26, 1995.

6. Morita, M.; Saito, S.; Shinohara, R.; Aoyagi, R.; Arita, M.; Kobayashi, Y. *Synlett* **2020**, *21*, 718–722.

#### Data for **5** ($^1$H NMR)

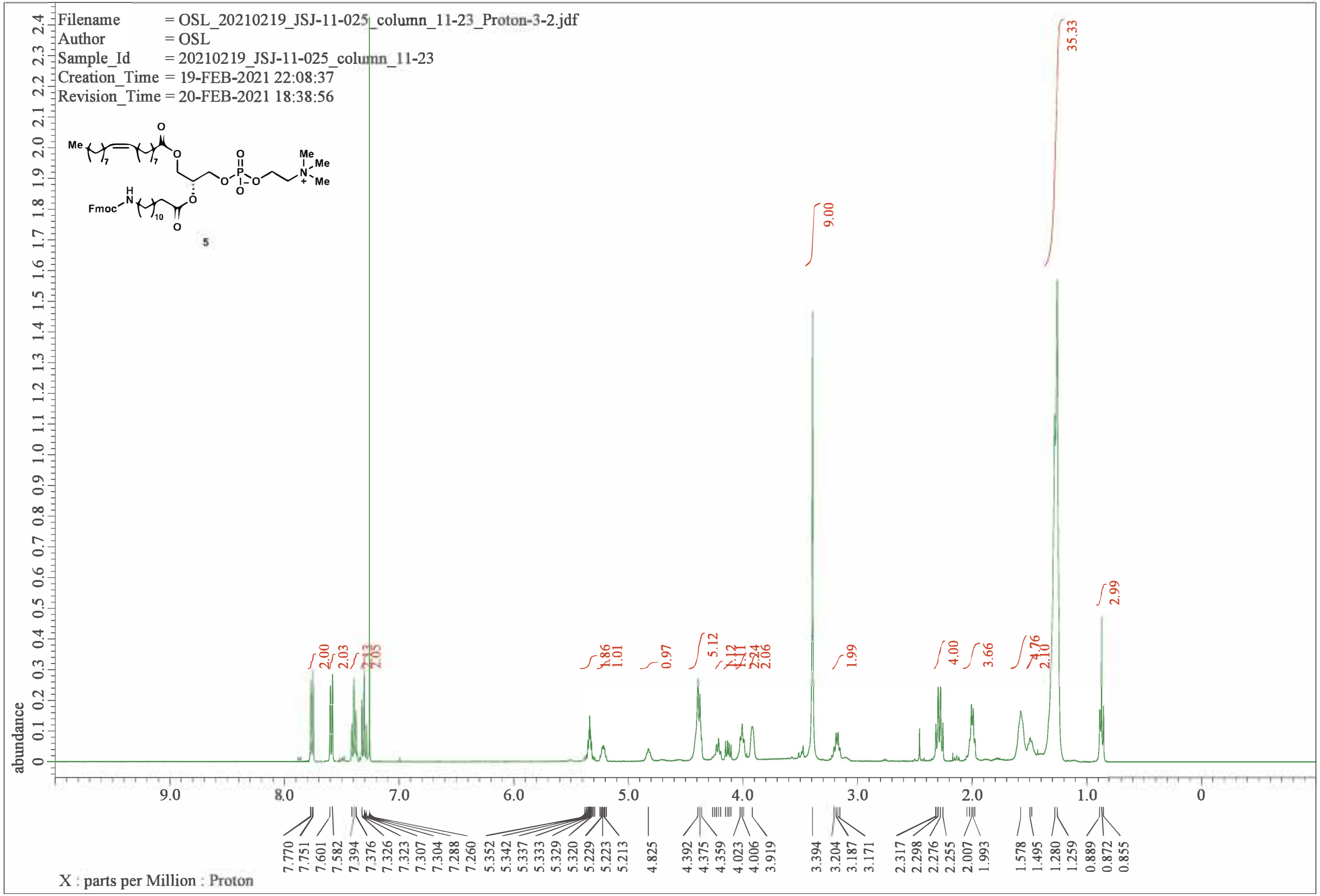

### Data for **5** ($^{13}$C NMR)

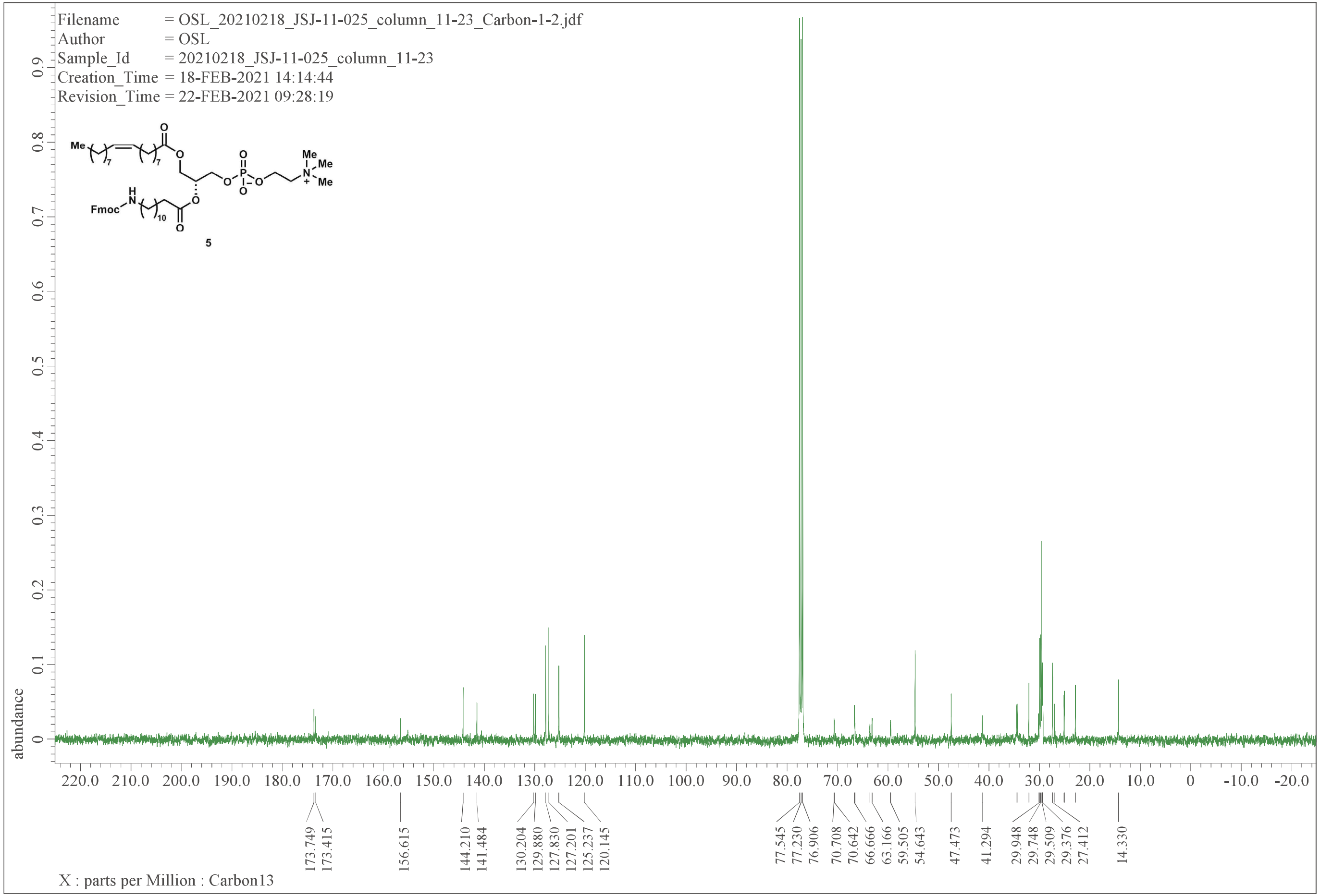

### Data for **5** ($^{31}P$ NMR)

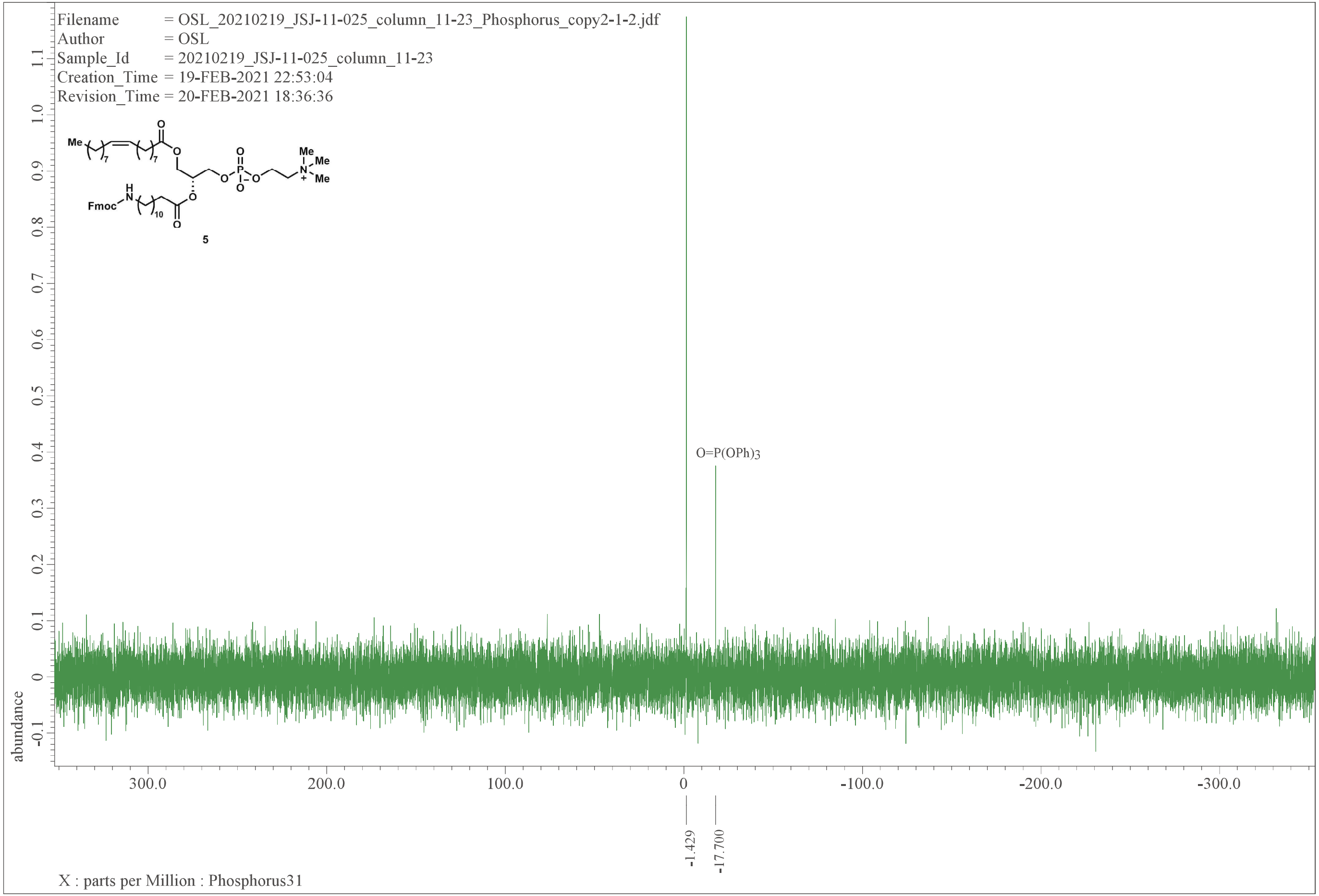

### Data for 6 ($^1$H NMR)

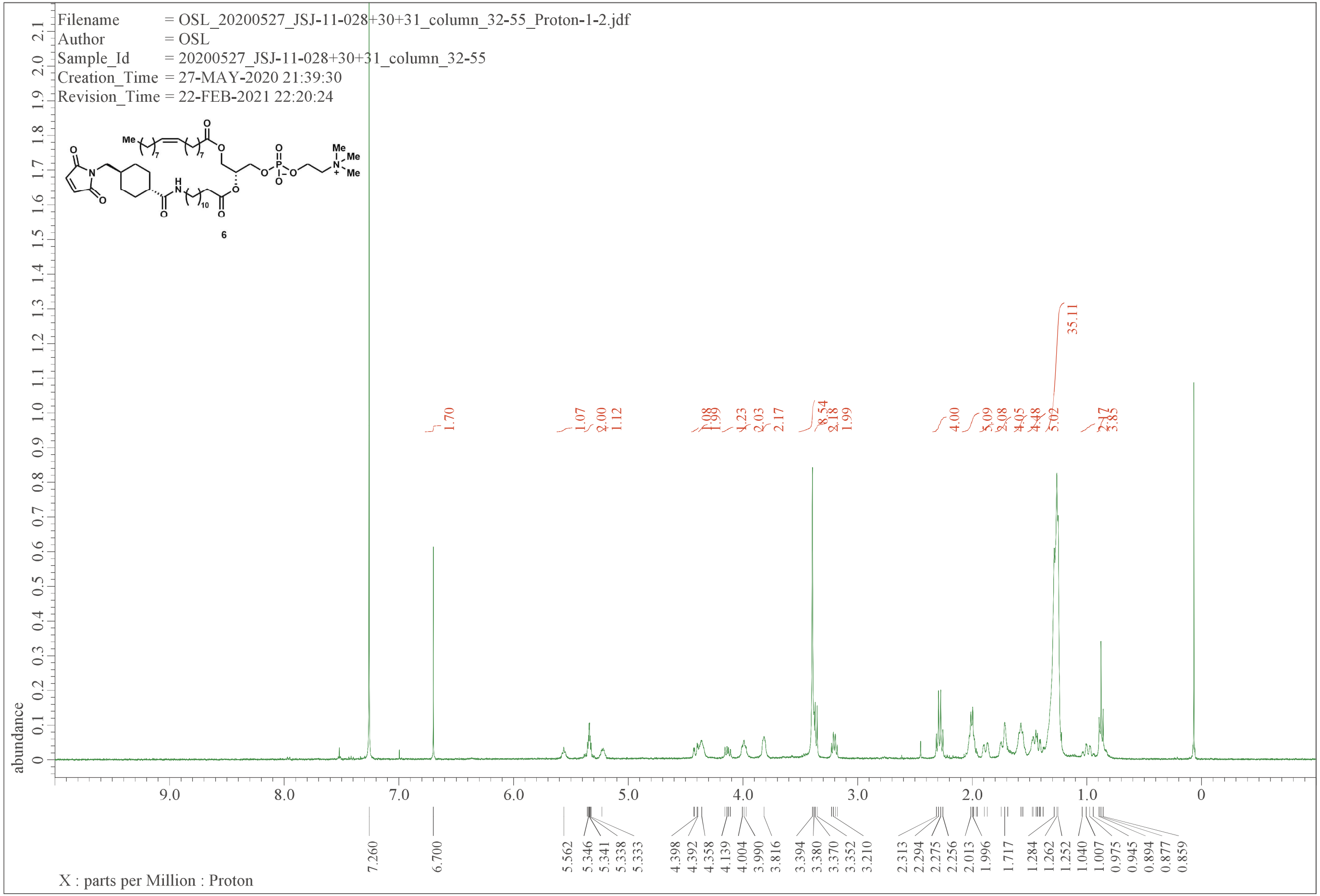

### Data for **6** ($^{13}$C NMR)

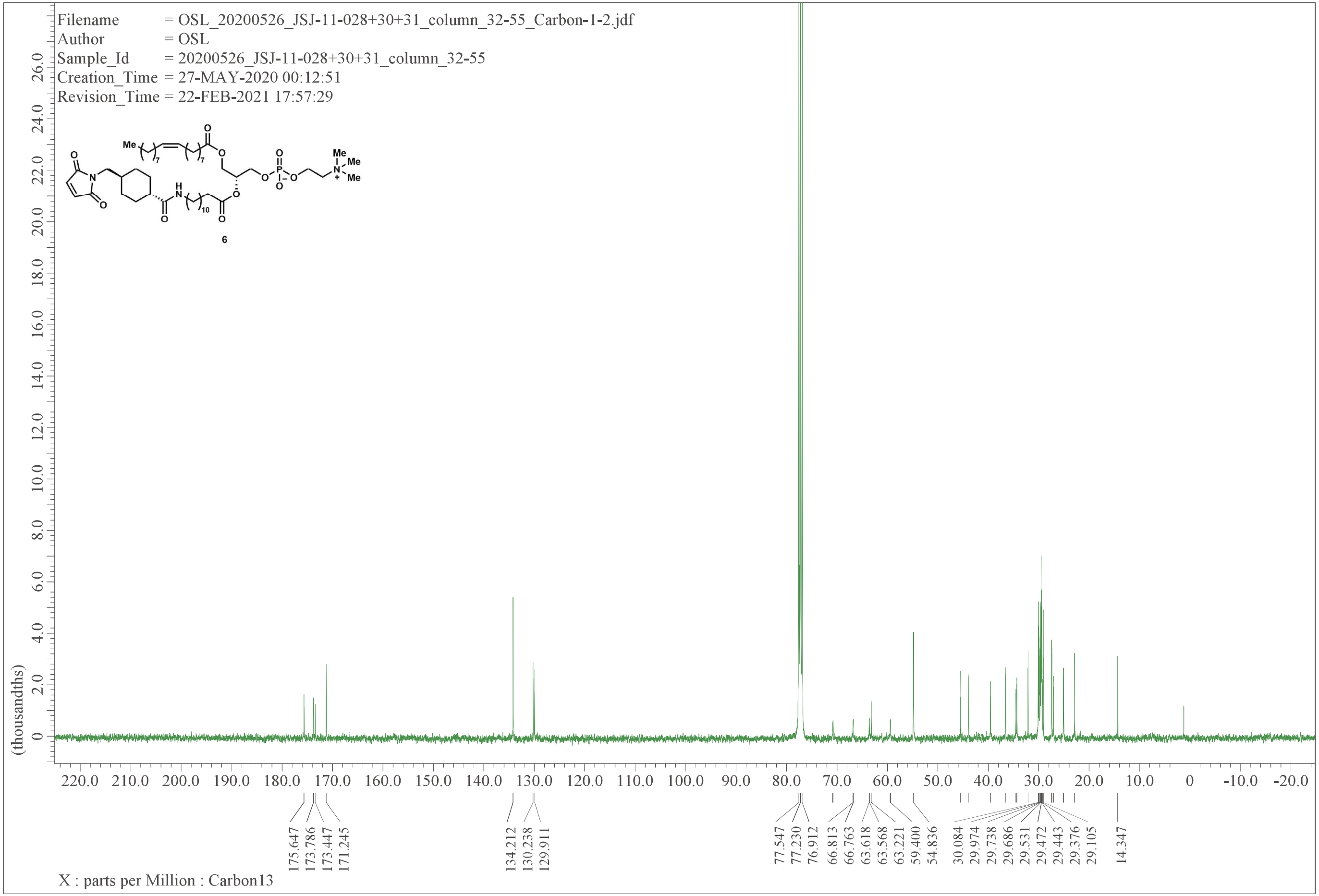

### Data for **6** ($^{31}$P NMR)

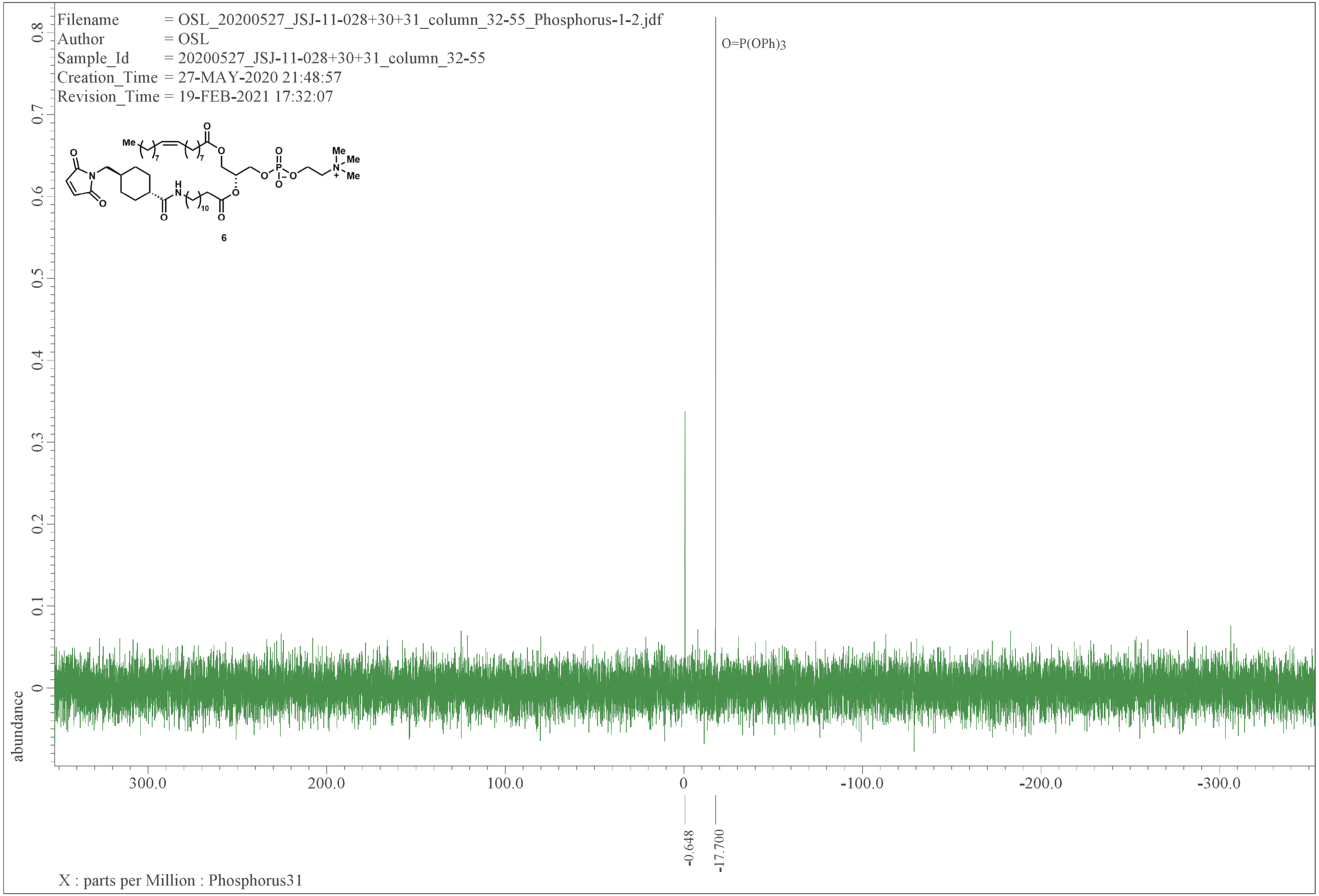
